# A Computational Perspective on the No-Strong-Loops Principle in Brain Networks

**DOI:** 10.1101/2025.09.24.678310

**Authors:** Fatemeh Hadaeghi, Kayson Fakhar, Moein Khajehnejad, Claus C. Hilgetag

## Abstract

Cerebral cortical networks in the mammalian brain exhibit a non-random organization in which reciprocal projections, although widespread, are systematically asymmetric in strength: feedforward connections are consistently stronger than their feedback counterparts, particularly in sensory cortices. This “no-strong-loops” principle is thought to prevent runaway excitation and maintain stability, yet its actual computational impact remains unclear. Here, we use computational analysis and modeling to show that connectivity asymmetry supports high working-memory capacity, whereas increasing reciprocity reduces memory capacity and representational diversity in reservoir-computing models of recurrent neural networks. We systematically examine synthetic architectures inspired by mammalian cortical connectivity and find that sparse, modular, and hierarchical networks achieve superior performance, relative to random, small-world, or core-periphery graphs, but only when reciprocity is constrained. Validated on directed mammalian (macaque, marmoset, rat, and mouse) connectomes, these results indicate that restricting reciprocal motifs yields functional benefits in sparse networks, consistent with an evolutionary strategy for stable, efficient information processing in the brain. These findings suggest a biologically-inspired design principle for artificial neural systems.

---

Biological neural networks have evolved under competing pressures to optimize energy costs of wiring and metabolism, while enhancing communication efficiency and computational capability. Such trade-offs have given rise to non-random architectural motifs that support efficient information processing under physical constraints (Van den Heuvel et al., 2016; Chen et al., 2006; Kaiser, 2007). One prominent, underexplored feature of cortical organization is the systematic asymmetry of reciprocal projections: while bidirectional connections between cortical areas are common, they are not balanced in strength, particularly in sensory cortices (Theodoni et al., 2022; Peng et al., 2024). This pattern is captured by the no-strong-loops hypothesis, which posits that, although reciprocal long-range connections are widespread in the cortex (Felleman and Van Essen, 1991), they follow a consistent structural asymmetry, with feedforward (FF) projections typically being stronger and exerting a driving influence, while feedback (FB) projections are weaker and modulatory. Moreover, the two types of projections are disentangled by targeting distinct cortical layers (Crick and Koch, 1998). This systematic avoidance of strong reciprocal motifs may imply a deeper functional constraint shaping cortical connectivity.

Experimental evidence supports the no-strong-loops hypothesis and demonstrates a systematic asymmetry in feedforward and feedback connection strengths across species. The connectional symmetry of the human brain is not directly accessible, as current noninvasive techniques for establishing brain connectivity cannot determine directionality in terms of structural connectivity (Kale et al., 2018). However, in non-human primate systems, such as the marmoset cortex, FF connections exhibit significantly higher mean strength than FB connections (Theodoni et al., 2022). In macaque visual cortex, FF pathways, though fewer in number, contribute comparable cumulative connection strength to the more numerous FB pathways, implying that individual FF connections are stronger on average (Markov et al., 2014). A similar pattern is observed in mouse visual cortex, where anterograde tracing shows higher FF than FB strength (D’Souza et al., 2022). These cross-species findings highlight a consistent pattern of asymmetric reciprocity in the organization of long-range cortical connections.

Despite its anatomical prominence, the computational role of asymmetric reciprocity and its gradients is still poorly understood. By contrast, connection features such as sparsity, modularity, and hierarchical organization have been linked to functional advantages in brain networks (Meunier et al., 2010; Hilgetag and Goulas, 2020; Hilgetag et al., 2000).

Reciprocity can be characterized in two distinct forms: structural (link) reciprocity, which refers to the tendency for pairs of neurons or regions to form mutual connections, and strength (weight) reciprocity, which captures the degree of symmetry in connection weights between reciprocally connected pairs. Both forms are evident in anatomical data (Theodoni et al., 2022; Liu et al., 2020; Rubinov et al., 2015), yet their functional implications remain unresolved, even though it is clear that reciprocity strongly shapes the dynamics and stability of recurrent systems (Shao and Ostojic, 2023; Dahmen et al., 2020; Peng et al., 2024). Therefore, a fundamental understanding of how symmetry features of reciprocity affect computation is critical for determining how they support the brain’s capacity for flexible, high-dimensional representations required for cognitive functions.

To address this gap, we systematically examine how reciprocal connectivity affects the computational capacity of recurrent neural systems. We use reservoir computing (RC) as a model framework (Jaeger and Haas, 2004) which provides a biologically inspired and computationally efficient approach that isolates the role of network structure from learning dynamics (Pascanu et al., 2013; Rodriguez et al., 2019; Damicelli et al., 2022; Suárez et al., 2024). This feature makes RC especially well-suited for assessing how architectural features shape computation in recurrent systems. In order to ensure the general applicability of our findings, we evaluate performance across a range of synthetic network topologies, including small-world, modular, hierarchical, and core-periphery architectures, each designed to reflect characteristic organizational patterns observed in mammalian connectomes. We further extend the analysis to empirical, directed connectomes derived from the long-distance neural pathways of the macaque brain (Young, 1993; Sporns and Zwi, 2004; Hilgetag et al., 2000), the macaque visual cortex (Markov et al., 2014), and the cortical connectivity of the common marmoset (Majka et al., 2016). Cross-species generalizability beyond the primate lineage is further examined using directed weighted connectomes of the rat (Bota et al., 2015) and mouse cortex (Rubinov et al., 2015). To manipulate reciprocity, we use two algorithms that we developed recently (Hadaeghi et al., 2026): the first adjusts structural (link) reciprocity in binary networks while preserving overall connection density, and the second one adjusts strength (weight) reciprocity in weighted networks while preserving the total sum of weights. We systematically vary network density, and for each density level, generate networks with graded amounts of reciprocity. This approach enables a controlled investigation of how both structural and strength reciprocity shape network computation across different architectural and sparsity regimes.

Our results show that increasing reciprocity consistently degrades both memory capacity and representational diversity across all network architectures and densities. This decline is observed not only in biologically inspired topologies but also in matched random networks that share the same density and reciprocity levels. When comparing network classes, we find that architectures resembling cortical organization outperform random counterparts under sparse conditions, but only when reciprocity is kept low. This performance advantage is most evident in networks with limited overall connectivity, where even moderate levels of reciprocity lead to significant degradation. In more moderately connected regimes, bio-inspired architectures continue to exhibit superior performance over random networks across a broader range of reciprocity values. However, overall computational capacity declines with increasing density across all architectures, suggesting that sparsity and constrained reciprocity jointly support optimal dynamical performance.

These findings, which we also validate for available directed empirical connectomes of non-human primates, highlight a fundamental trade-off between reciprocal connectivity and computational efficiency. In biological systems, this trade-off may explain the evolutionary suppression of strong loops. In artificial systems, it provides a design principle for developing energy-efficient, sparsely connected recurrent networks. The robustness of these conclusions is further confirmed across nonlinear tasks, application to additional species beyond the primate lineage, and larger network scales. By linking a key anatomical constraint of cortical organization, namely, limited reciprocity, to its computational consequences, our work offers a principled explanation for the suppression of strong reciprocal motifs in the brain.

## 2 Results

### 2.1 Overview of the computational framework

To systematically evaluate how reciprocity affects recurrent computation, we adopted a reservoir computing (RC) framework in which bio-inspired and empirical networks serve as the recurrent core. Across experiments, we manipulated two forms of reciprocity, structural (link) and strength (weight) reciprocity, and evaluated network performance using two key metrics: memory capacity (temporal information retention) and kernel rank (representational diversity). Crucially, both metrics are task-agnostic: they reflect intrinsic computational capacity and representational resources of the network, independent of any specific input-output mapping. This makes them general measures of how network structure shapes the computational regime, rather than proxies for performance on a particular task. An overview of the workflow is shown in Figure 1.

**Figure 1.**
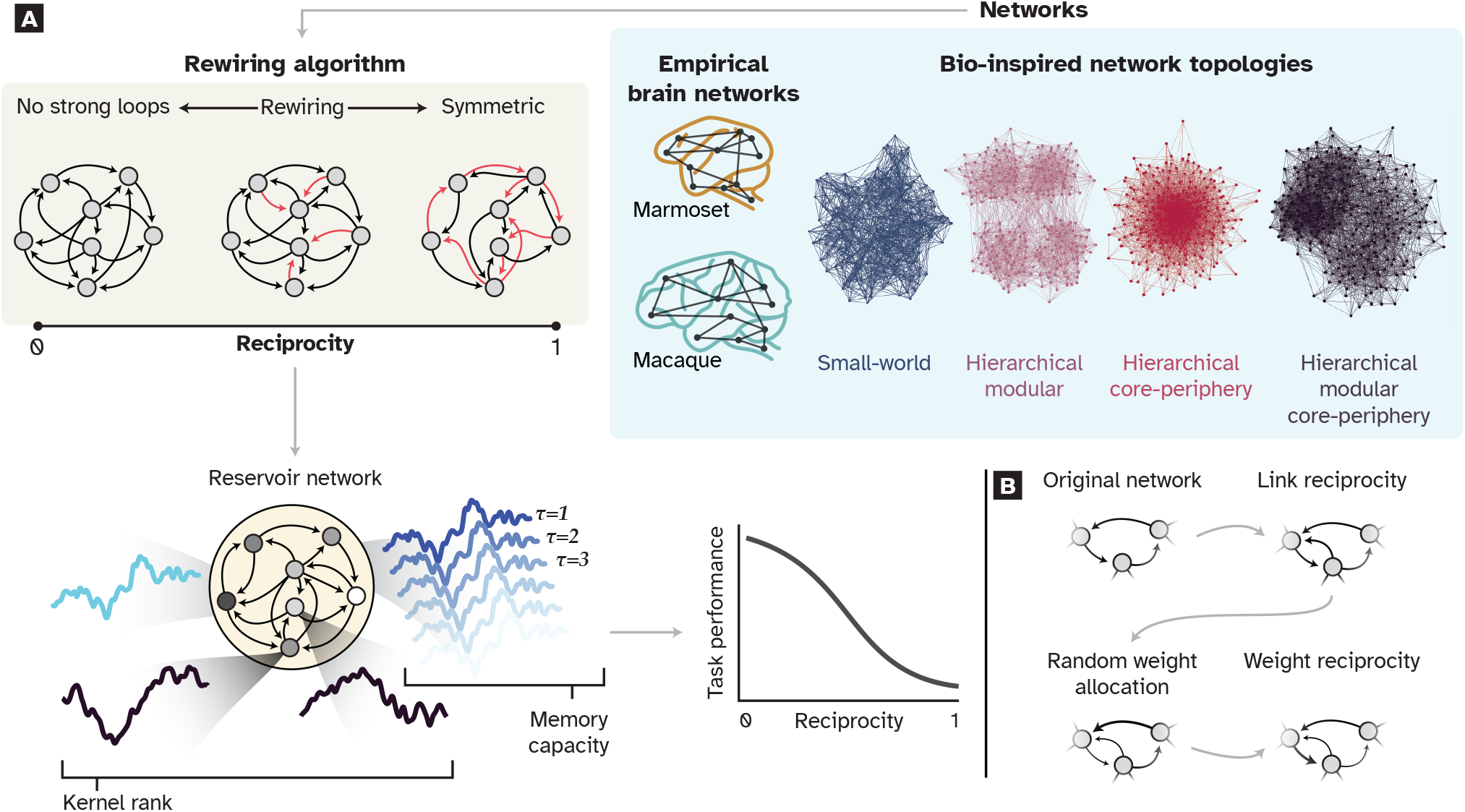
Overview of the computational framework and experimental design. (**A**) Networks with varying topologies and reciprocity levels served as the recurrent cores of reservoir computing models. *Top right:* Two classes of networks were used: empirical directed connectomes of macaque and marmoset cortex, and four classes of synthetic bio-inspired topologies (small-world, hierarchical modular, hierarchical core-periphery, and hierarchical modular core-periphery). *Top left:* Reciprocity was systematically varied from fully asymmetric (no strong loops) to fully symmetric using the network reciprocity control algorithm (Hadaeghi et al., 2026), illustrated here for link reciprocity: unidirectional connections (black arrows) are progressively converted into bidirectional pairs (red arrows) while preserving connection density. *Bottom left:* Each network served as the fixed recurrent reservoir of a leaky-integrator echo state network, receiving a continuous input stream and projecting to a trainable linear readout. Network performance was quantified using two task-agnostic metrics: memory capacity (MC), measuring the ability to linearly reconstruct past inputs across increasing time lags (τ = 1, 2, 3, …), and kernel rank (KR), measuring the dimensionality of the internal representation space. *Bottom right:* Across all conditions, increasing reciprocity produced a consistent decline in task performance. (**B**) The two forms of reciprocity were manipulated in this study. *Link reciprocity* (top row) refers to the proportion of bidirectional connections in the binary network structure: an asymmetric network with predominantly unidirectional edges is rewired so that connections become mutually reciprocal. *Strength reciprocity* (bottom row) refers to the symmetry of connection weights between reciprocally connected node pairs: weights are initially assigned independently from a log-normal distribution (random weight allocation), yielding asymmetric strengths even when both directions of a connection exist; strength reciprocity is then enforced by adjusting weights so that each reciprocal pair becomes symmetric. These two forms of reciprocity are manipulated independently, allowing their computational consequences to be dissociated.

### 2.2 Network architectures and reciprocity manipulation

To investigate how cortical wiring constraints influence computational performance, we first generated synthetic networks that capture key topological features of mammalian connectomes. Synthetic models allow systematic control of reciprocity while avoiding limitations of empirical data, such as noise, incomplete coverage, and inter-individual variability. We then validated reciprocity effects observed in synthetic networks using directed connectomes of macaque, marmoset, rat, and mouse cortex, ensuring that the computational effects of reciprocity observed in synthetic networks also hold in real brain networks.

In addition to small-world (SW) (Watts and Strogatz, 1998; Fakhar et al., 2022) and hierarchical modular (HM) networks (Ravasz et al., 2002; Rodriguez et al., 2019; Milisav et al., 2025), we introduced two novel classes of brain-inspired networks: hierarchical core-periphery (HCP) and hierarchical modular core-periphery (HMCP). Core-periphery organization, with a dense core and sparse periphery, is thought to support robust and flexible communication (Borgatti and Everett, 2000), yet its computational role under varying reciprocity remains unexplored.

Network generation and reciprocity manipulation are detailed in Materials and Methods.

### 2.3 Spectral properties reveal topology-dependent impact of reciprocity

Spectral properties are central to recurrent network dynamics. We examined how reciprocity affects three key measures: spectral radius, spectral gap, and eigenvalue non-normality. The spectral radius, the largest magnitude among the eigenvalues of the weight matrix, provides the exact asymptotic decay or growth rate of network states over time in linear systems. When the spectral radius *ρ* < 1, past inputs decay exponentially; as *ρ* approaches 1.0, memory length increases, and values exceeding 1.0 cause unbounded growth. In nonlinear reservoirs, saturation prevents explosion, but optimal computation still requires *ρ* near 1.0, as a radius that is too low drives activity toward zero, while one that is too high (particularly *ρ* > 1) compresses dynamic range and reduces input sensitivity. In the present setting, the spectral radius rose steadily with reciprocity across all architectures, driven by a dominant positive real eigenvalue, pushing networks away from this optimal regime. All networks were therefore normalized to maintain *ρ* below 1.0, following reservoir computing practice (Jaeger and Haas, 2004; Farkaš et al., 2016).

The spectral gap, defined as the difference between the largest and second-largest eigenvalue magnitudes, determines how quickly activity collapses onto the dominant eigenmode. A large gap causes rapid convergence to a single direction, compressing the representational space; a small gap allows multiple eigenmodes to persist simultaneously, supporting richer internal dynamics. Here, reciprocity reduced the gap across architectures, narrowing the dynamical range. Declines were roughly linear in small-world, core-periphery, and random networks (null models), but slower and nonlinear in modular ones (HM, HMCP), suggesting greater robustness.

Eigenvalue non-normality, measured by Henrici’s index (detailed in Spectral property analysis), quantifies the departure from orthogonality of eigenmodes. High non-normality enables strong transient amplification even when all eigenvalues are stable, allowing the network to represent inputs along many directions simultaneously and thereby supporting longer memory and richer dynamics. At low reciprocity, networks showed high non-normality, reflecting directional asymmetry. With increasing reciprocity, eigenvalue distributions became more symmetric and Henrici’s index declined, reducing transient diversity — consistent with prior findings on links between reciprocity and spectral symmetry (Asllani et al., 2018; Hennequin et al., 2012).

Figure 2 and Supplementary Figures S1 and S2 summarize how spectral radius, spectral gap, and Henrici’s index change as a function of link reciprocity across network types, sizes, and sparsity levels. Together, these results show that reciprocity systematically alters spectral structure, with modular architectures exhibiting greater resilience to these changes. The implications for computation are elaborated in subsequent sections.

**Figure 2.**
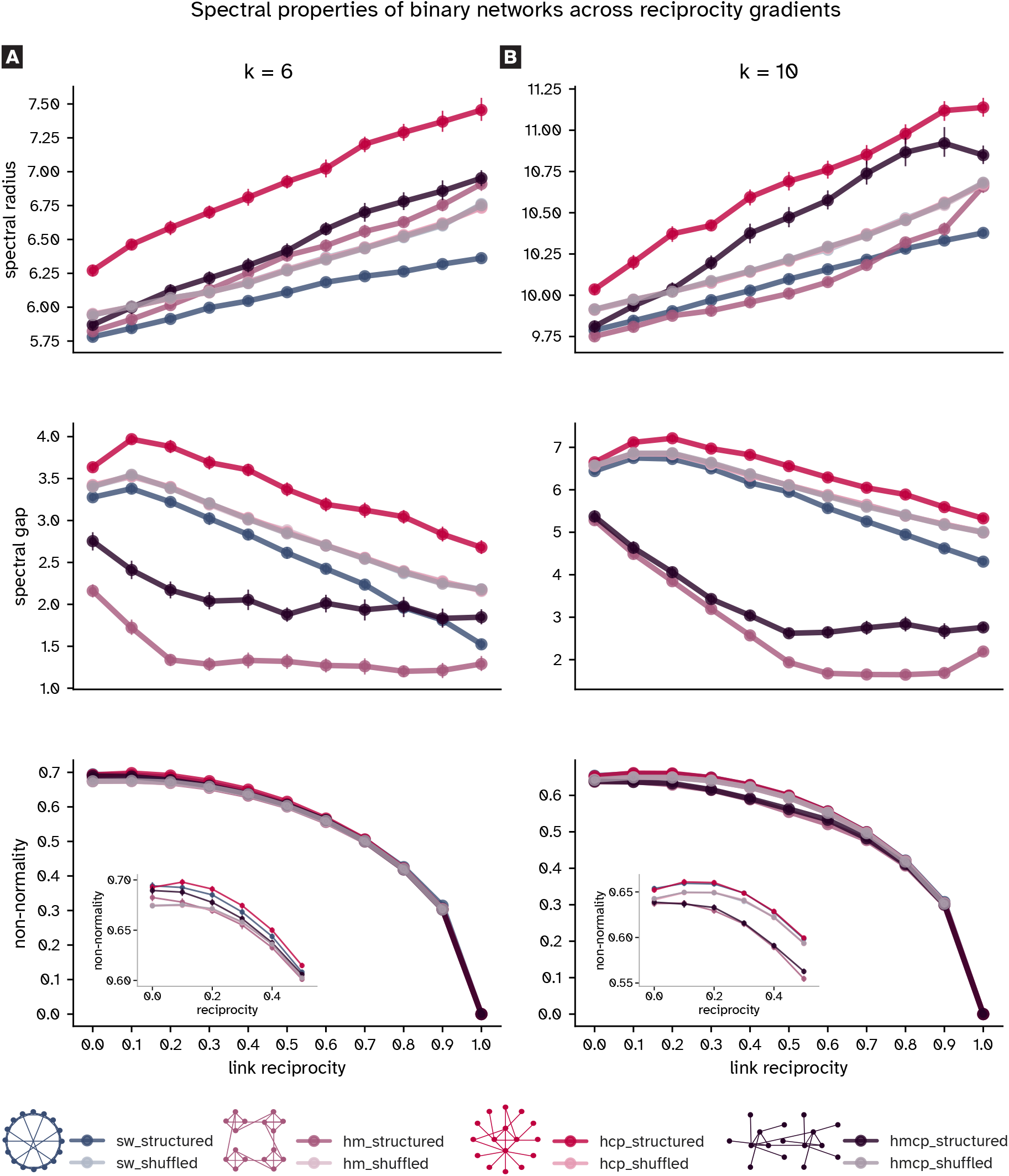
Spectral properties of synthetic networks across reciprocity gradients. Shown are the spectral radius, spectral gap, and Henrici’s index of non-normality for binary networks with 64 nodes under two connectivity levels: (**A**) lower connectivity (mean out-degree *k* = 6) and (**B**) higher connectivity (mean out-degree *k* = 10). Each point represents the mean across 50 instances; error bars denote standard deviation. In most conditions, variability across network instances was low. The spectral radius increased monotonically with link reciprocity across all architectures, while the spectral gap narrowed and non-normality declined. Insets in the non-normality panels show an expanded view of the low reciprocity range (0–0.5), where architecture-dependent differences are most pronounced before convergence at higher reciprocity levels. Together, these results illustrate how increasing reciprocity systematically restructures the spectral properties of diverse network architectures.

### 2.4 Link reciprocity reduces memory and representational capacity across network architectures

To investigate the impact of reciprocity on computational capacity, we conducted a memory task on reservoirs of varying sizes (*N* = 64, 128, and 256 nodes), spanning the range of cortical parcellations commonly used in connectomics. We also assessed representational diversity using the kernel rank (KR) metric (Legenstein and Maass, 2007; Dale et al., 2021), which quantifies the separation property of network dynamics. Both measures are described in Materials and Methods. For each network type, we generated 50 trials at two mean out-degrees: *k* = 6 (lower connectivity) and *k* = 10 (higher connectivity). The spectral radius and other network parameters were optimized to maximize memory capacity.

Across all experiments, reciprocity reduced both memory capacity and kernel rank (Figure 3). Modularity consistently enhanced performance, explaining why HMCP networks outperformed core–periphery architectures lacking modules. Increasing network size at fixed mean out-degree produced sparser networks, which tended to support higher memory and representational diversity. Still, across sizes and architectures, both metrics declined with increasing reciprocity (Supplementary Figure S3). For the representative case of HM networks at *k* = 6 and N = 64, MC declined linearly with reciprocity (slope = − 1.26, *R*^2^ = 0.995, *P* < 0.001) and KR showed a consistent negative trend (slope = − 0.45, *R*^2^ = 0.882, *P* < 0.001). Complete mean performance values and structured-vs-null comparisons across all experimental conditions are reported in Supplementary Table S1. In addition, slopes for all network types, sizes, and connectivity regimes are reported in Supplementary Table S2. For consistency, we applied network generation parameters optimized for 64-node networks across all sizes. To confirm robustness, we also re-optimized parameters for 128- and 256-node networks and again observed declines in both metrics (Supplementary Figure S4). Subsequent analyses used non-optimized parameters to ensure consistency and avoid tuning-related variability.

**Figure 3.**
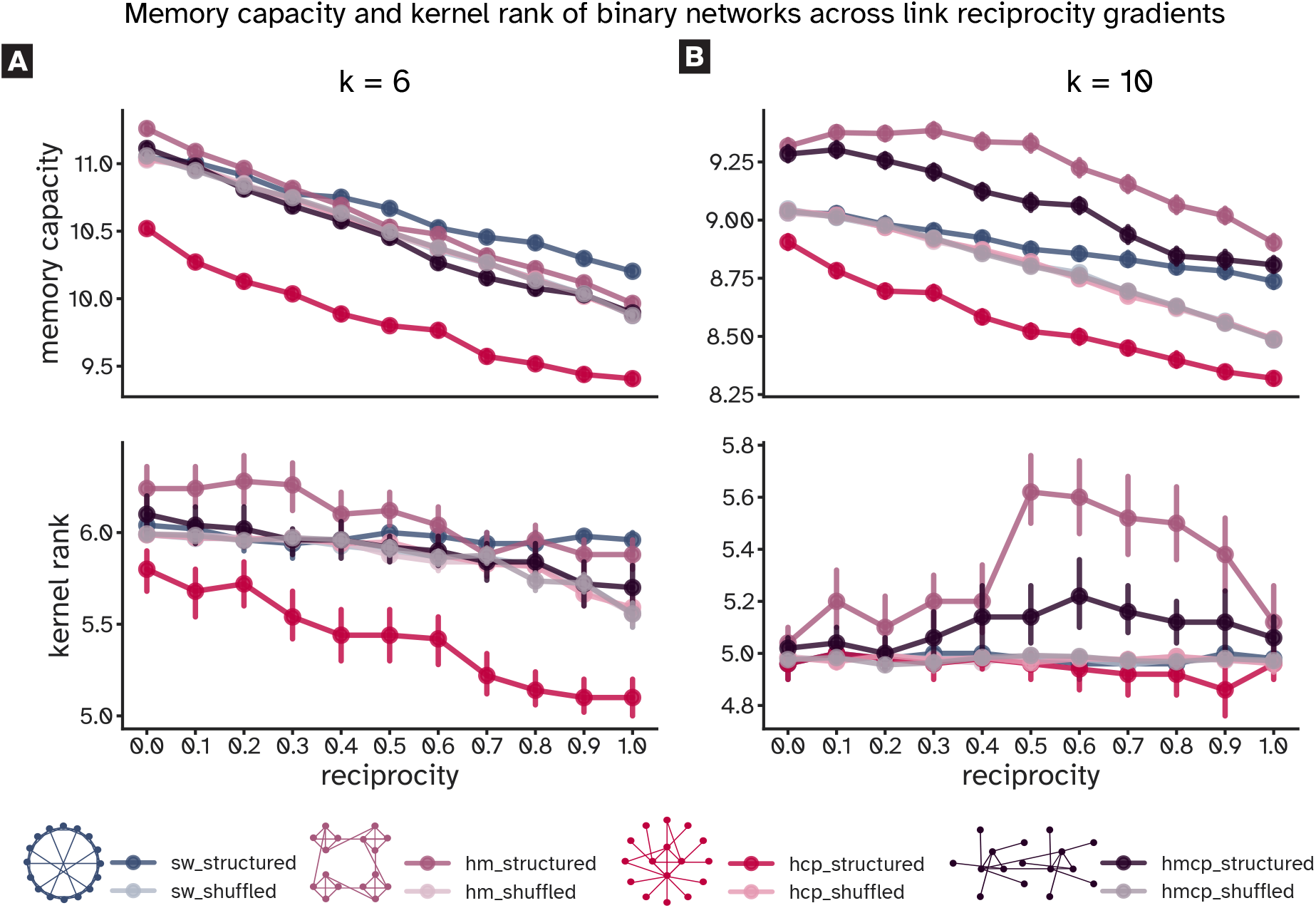
Symmetrical reciprocity reduces memory capacity and kernel rank in recurrent networks. Results are shown for binary networks with 64 nodes under (**A**) lower connectivity (mean out-degree *k* = 6) and (**B**) higher connectivity (mean out-degree *k* = 10). Error bars indicate standard deviation across 50 realizations. Across all experiments, increasing reciprocity led to a decline in memory capacity and kernel rank. This degradation was particularly pronounced in the lower connectivity regime (MC slope = {™0.86, −1.26, −1.07, −1.23} for {SW, HM, HCP, HMCP}, all *P* < 0.001; KR slope = {™0.09, −0.45, −0.76, −0.41}, SW: *P* = 0.031; HM, HCP, HMCP: all *P* < 0.001; Supplementary Table S2). The SW kernel rank slope, while statistically significant, is small in absolute magnitude and reflects a weak trend; the decline in kernel rank is most pronounced in HCP networks and most consistent across conditions in HM networks. In the memory task, HM networks significantly outperformed their random counterparts (*P* < 0.001; Cohen’s *d* from 0.18 at link reciprocity = 0.5 to 0.84 at reciprocity = 0.0). Small-world networks outperformed random networks at higher reciprocity values, with the gap between structured and shuffled SW networks widening progressively beyond reciprocity = 0.4 (Cohen’s *d* from 0.46 to 1.18, all *P* < 0.001). In the higher connectivity regime (**B**), MC slopes were shallower across all architectures (SW = −0.30, HM = −0.45, HCP = −0.56, HMCP = −0.56, all *P* < 0.001), with modular networks consistently outperforming random versions (*P* < 0.001, Cohen’s *d* ≥ 0.9). HM networks maintained a consistently higher kernel rank across all reciprocity values, with a peak at reciprocity = 0.5. Full slope statistics across all sizes are reported in Supplementary Table S2.

Modularity emerged as a key factor supporting diverse representations in both sparsity regimes. In lower connectivity networks, increasing reciprocity may tend to convert asymmetric inter-modular connections into reciprocal intra-modular ones, effectively increasing modularity (Hadaeghi et al., 2026) and reducing the diversity of inter-modular communication, which is consistent with the observed reduction in kernel rank (Supplementary Table S1, HM condition). In the higher connectivity regime, the greater number of connections, including long-range links, allowed pruning to balance intra- and inter-modular communication more effectively. This balance promoted more diverse representations, with reciprocity values around 0.5 yielding particularly high kernel rank in hierarchical modular networks. These divergent trends across sparsity levels highlight the joint influence of density and reciprocity in shaping the computational landscape of recurrent architectures.

### 2.5 Reciprocity effects persist in networks with heterogeneous weights

To test whether reciprocity-driven declines extend beyond binary connectivity, we assigned log-normally distributed weights to the same binary backbones. This introduced biologically realistic heterogeneity while preserving topology. As in the binary case, both memory capacity and kernel rank (KR) declined systematically with increasing reciprocity (Figure 4; Supplementary Figures S5, S6, S7, S8, and S9).

**Figure 4.**
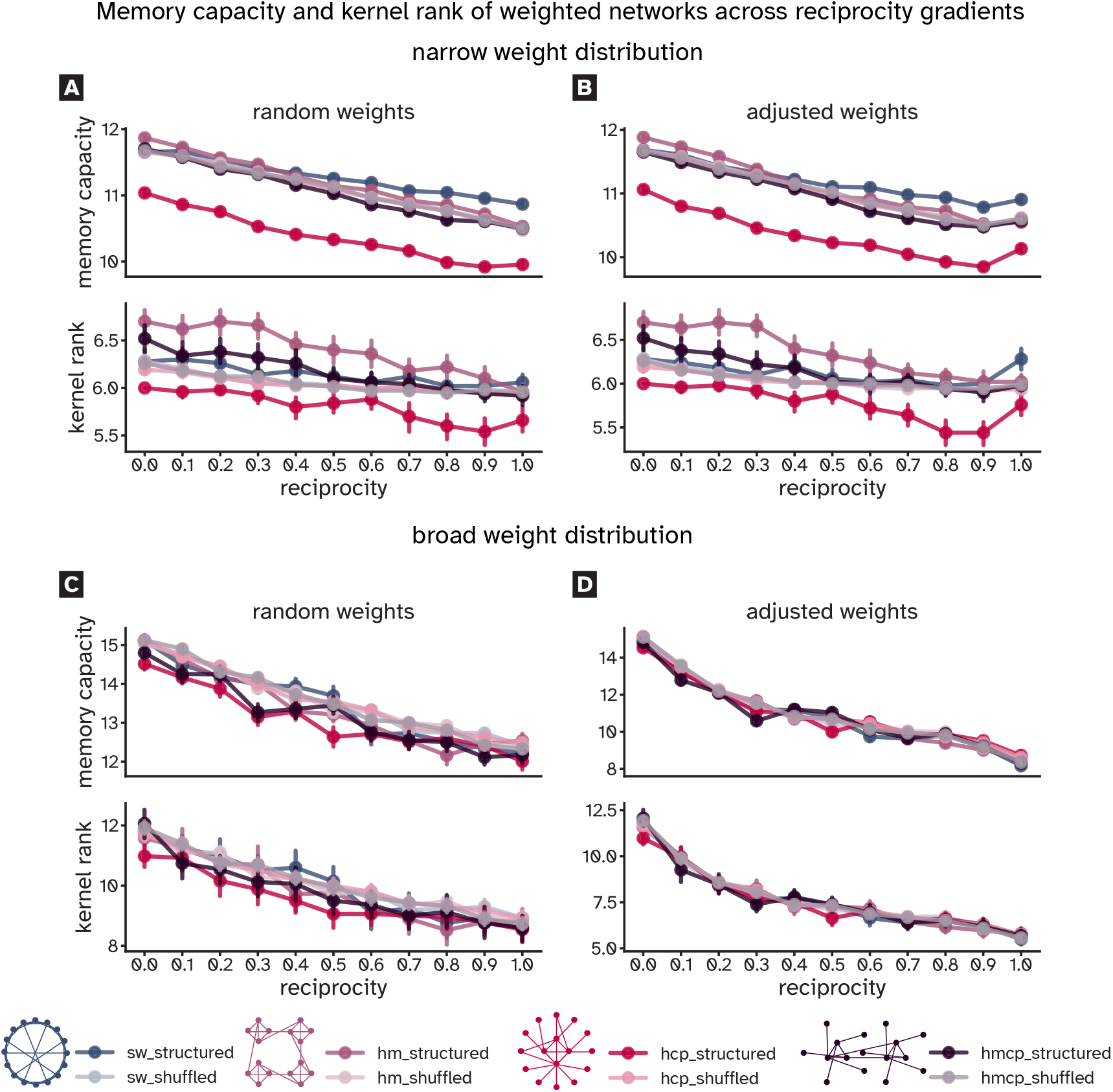
Impact of reciprocity on memory and kernel rank in weighted networks with random and adjusted strength reciprocity. Results are shown for 64-node networks with weights sampled from log-normal distributions, under lower connectivity (*k* = 6). Error bars indicate standard deviation across 50 trials. Panels **A** and **B** show results under narrow weight distributions (SD = 0.1, mean = 0.25); panels **C** and **D** show results under broad weight distributions (SD = 0.9, mean = 0.25). Left panels (**A, C**) show random weight allocation; right panels (**B, D**) show enforced strength reciprocity. Across both weight regimes, increasing link reciprocity leads to systematic declines in MC and KR. Under narrow weights and random allocation (**A**), MC declined linearly (MC slope: SW = −0.82, HM = −1.29, HCP = −1.12, HMCP = −1.24, all *P* < 0.001; KR slope: SW = − 0.31, HM = −0.62, HCP = −0.52, HMCP = −0.54, all *P* < 0.001). Under broad weights and random allocation (**C**), declines were steeper (MC slope: SW = −2.89, HM = −2.79, HCP = −2.30, HMCP = −2.63, all *P* < 0.001; KR slope: SW = −3.15, HM = −3.02, HCP = −2.39, HMCP = −2.86, all *P* < 0.001). Under random weight assignment, strength reciprocity remained largely independent of link reciprocity (0.4 ± 0.25 and 0.1 ± 0.08 for SD = 0.1 and 0.9, respectively). When strength reciprocity was explicitly enforced (**B, D**), declines were sharper. Under narrow weights (**B**), declines remained linear (MC slope: SW = −0.86, HM = −1.41, HCP = −1.07, HMCP = −1.23, all *P* < 0.001). Under broad weights (**D**), declines were exponential (*R*^2^ ≥ 0.93), with MC decay rates of SW = −2.14, HM =™ 2.16, HCP = −2.64, HMCP = −2.21 (all *P* ≤0.05) and KR decay rates of SW = −3.19, HM = −3.03, HCP = −3.28, HMCP = −3.89 (all *P* 0.01), indicating that representational diversity is particularly sensitive to enforced weight symmetry. Full statistics are reported in Supplementary Tables S1 and S2.

This effect was robust across a range of weight distributions. In particular, we tested both narrow and broad standard deviations of the log-normal distribution and found that increasing reciprocity reduced memory and representational capacity in all cases. The decline was steeper when weights were drawn from longer-tailed distributions, which introduced greater variability and stronger connections. For HM networks at *k* = 6 and N = 64, MC declined linearly with reciprocity under both weight regimes (narrow: slope = −1.29, *R*^2^ = 0.994, *P* < 0.001; broad: slope = −2.79, *R*^2^ = 0.906, *P* < 0.001), confirming that broader weight distributions steepen rather than attenuate the reciprocity-driven decline. Full statistics across all conditions are reported in Supplementary Table S2.

Although broader weight distributions improved absolute performance, they did not eliminate reciprocity’s negative impact. Under these conditions, performance differences between random and bio-inspired architectures were less pronounced, with most networks performing comparably across reciprocity levels and only a few showing notable memory differences. Hierarchical core-periphery (HCP) networks performed on par with others, overcoming the gap seen in binary experiments (Figure 4, panels **A** and **C**; Supplementary Figures S5, S6, S7, S8, and S9). HCP networks also showed a shallower memory decline (slope ≈ −0.23) with increasing reciprocity, suggesting greater resilience to reciprocal connectivity (Figure 4, panel **C**).

These results show that while weight heterogeneity enhances absolute performance, it does not buffer recurrent networks from the computational costs of reciprocity. Even in networks with biologically plausible weight distributions, reciprocity remains a dominant factor limiting memory and representational diversity.

### 2.6 Enforced strength reciprocity exponentially degrades memory and representation quality

To directly assess how strength reciprocity affects computation, we applied our previously developed algorithm (Hadaeghi et al., 2026) (see Section Materials and Methods) to enforce reciprocal weight symmetry. Log-normal weights, initially assigned independently, were adjusted so that strength reciprocity matched link reciprocity. This procedure isolated the specific impact of reciprocal weight symmetry on memory and representation quality.

Enforcing strength reciprocity produced a near-exponential decline in both memory capacity and kernel rank across architectures and network sizes (Figure 4, panels **B** and **D**; Supplementary Figures S5, S6, S7, S8, and S9). This degradation was especially pronounced when weights were sampled from long-tailed log-normal distributions, which introduce greater heterogeneity. In these conditions, reciprocal weight symmetry severely impaired the network’s ability to retain temporal information and generate diverse internal states.

With random weight assignments, strength reciprocity values remained low and largely independent of link reciprocity (Figure 4 caption). In contrast, when reciprocity was explicitly enforced, performance deteriorated sharply, with exponential fits capturing the decline across all conditions. For HM networks at *k* = 6 and N = 64 with broad log-normal weights, the fitted exponential decay rates were 2.16 for MC (*R*^2^ = 0.980, *P* < 0.001) and 3.03 for KR (*R*^2^ = 0.974, *P* < 0.001), with KR exhibiting a steeper decline than MC, indicating that representational diversity is particularly sensitive to enforced weight symmetry. Full decay rates and fit statistics across all network types and conditions are reported in Supplementary Table S2.

These results indicate that enforcing strength reciprocity not only preserves, but also amplifies the computational decline associated with increasing reciprocity. Slope comparisons between random and enforced strength reciprocity conditions confirmed that the exponential decline under enforced reciprocity is significantly steeper than the linear decline under random weight assignment across all network types (Supplementary Table S2). While weight heterogeneity alone can boost absolute performance (as shown in the previous section), it fails to counteract the negative effects of reciprocal weight symmetry.

Comprehensive statistical results across all network types (SW, HM, HCP, HMCP), network sizes, connectivity regimes (*k* = 6, 10), and weight conditions (binary, narrow and broad random weights, narrow and broad enforced weights), spanning 2,200 experimental conditions in total, are reported in Supplementary Table S1.

### 2.7 Spectral properties explain performance degradation with increasing reciprocity

We analyzed how reciprocity shapes recurrent network dynamics by relating spectral properties to memory capacity and kernel rank. As reciprocity increased, via structural links or enforced weight symmetry, the spectral radius rose across architectures, driven by a dominant positive real eigenvalue that pushed networks toward instability (Figure 2; Supplementary Figures S1 and S2).

To ensure stable comparisons across conditions, we applied spectral normalization to all networks, tuning them to operate near the critical regime required to satisfy the echo-state property (Jaeger and Haas, 2004; Farkaš et al., 2016)in reservoir computing. After normalization, spectral radii were close to 1.0 across architectures, reciprocity levels, sizes, and sparsities. High-reciprocity networks, however, required stronger weight suppression to offset the elevated radius from reciprocal motifs, substantially reducing total connection strength and recurrent gain. Without normalization, MC was uniformly poor across all reciprocity levels and connectivity regimes (Supplementary Figure S10), confirming that networks must operate near the edge of stability for meaningful computation. The reciprocity-driven decline observed under normalization is therefore not an artifact of the procedure itself, but reflects the structural consequences of reciprocity within the optimal dynamical regime — a conclusion further supported by the gain compensation experiment below.

To test whether gain reduction is the sole mechanism underlying the MC decline, we compensated for it by scaling input weights *W*_in_ by the raw spectral radius *ρ* of each network. Despite this compensation, MC declined monotonically with reciprocity across all connectivity levels (all *R*^2^ ≥ 0.708, *P* ≤0.001; Supplementary Figure S10, with slopes at *k* = 5 and *k* = 10 that were 1.16 × and 1.49 × steeper than the uncompensated counterparts, respectively. The steeper decline under input scaling is consistent with the reservoir operating further from its optimal input drive regime: scaling *W*_*i*_ *n* upward increases the effective input magnitude received by each node, which, combined with the reduced recurrent gain imposed by spectral normalization on high-reciprocity networks, pushes the reservoir into a saturated or over-driven state that further degrades temporal memory. This result confirms that gain reduction is a genuine contributing factor but is insufficient to explain the performance decline, which additionally reflects eigenspectral restructuring, specifically the narrowing spectral gap and reduced non-normality, that persists independently of input gain. These two mechanisms are therefore dissociable and additive in their effects on memory capacity. Formal correlations confirm this dissociation: spectral radius was the dominant predictor of MC across all network types and reciprocity levels (*r* = −0.977, *P* < 0.001; Figure 5, panel **A**, and Supplementary Figure S11, Panel A). To be more clear, these results identify two distinct contributing mechanisms for the MC decline: gain reduction from aggressive spectral normalization, and eigenspectral restructuring that persists independently of gain.

**Figure 5.**
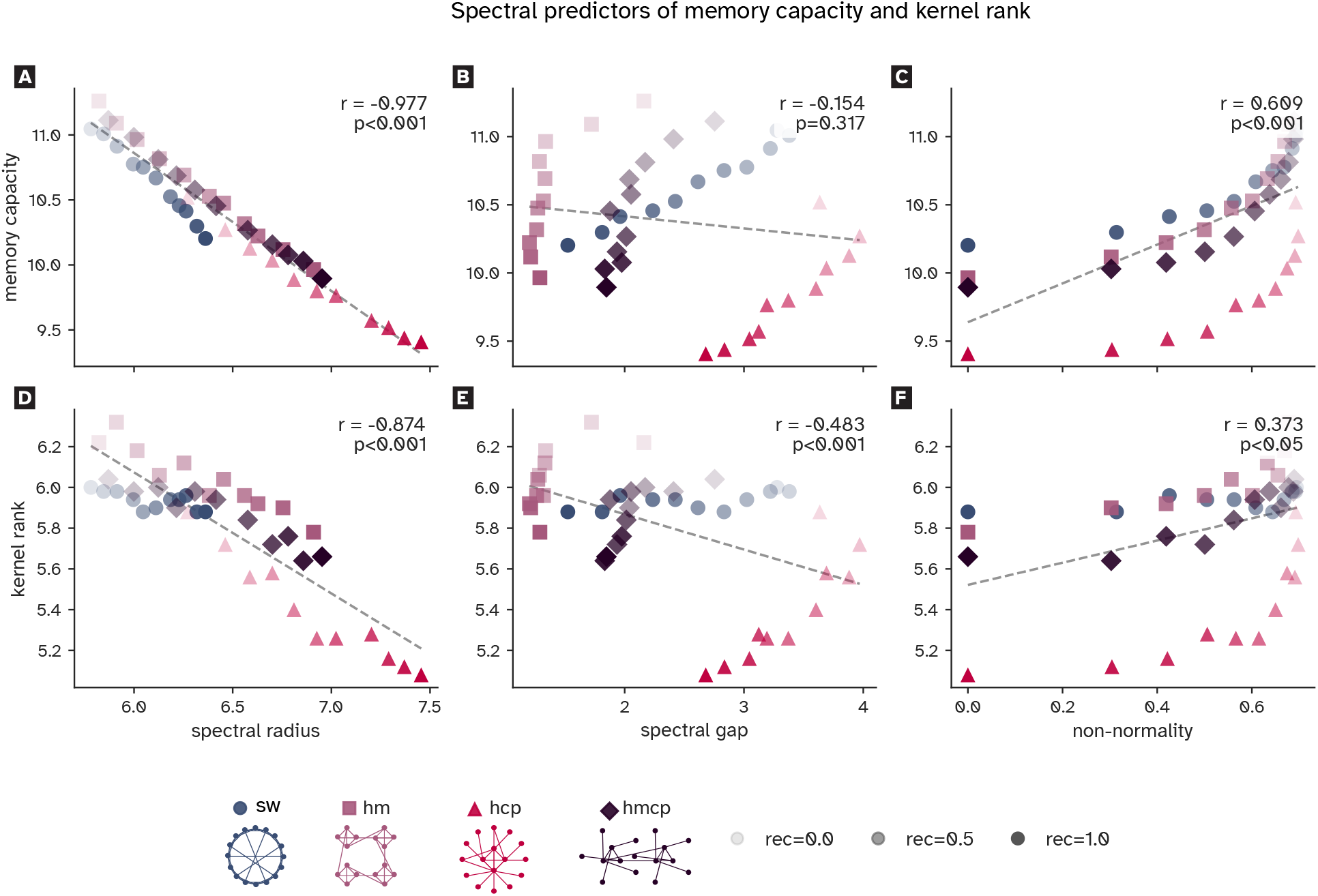
Spectral properties predict memory capacity and kernel rank across network architectures and reciprocity levels. Scatter plots show the relationship between three spectral properties (spectral radius, spectral gap, and non-normality) and two computational performance metrics (memory capacity, MC; kernel rank, KR) for binary networks with 64 nodes and *k* = 6 (see Supplementary Figure S11 for networks with *k* = 10). Each point represents the mean over 50 network realizations for one (network type × reciprocity) combination. Network type is encoded by colour and marker shape; reciprocity is encoded by marker opacity (low reciprocity = faint, high reciprocity = opaque). Dashed lines indicate linear regression fits across all structured networks combined. Pearson correlation coefficients (*r*) and *P*-values are reported for the pooled sample in each panel. Spectral radius (**A, D**) was strongly negatively correlated with MC (pooled: *r* = −0.977, *P* < 0.001; per network type: SW *r* = −0.994, HM *r* = −0.998, HCP *r* = −0.994, HMCP *r* = −0.998, all *P* < 0.001) and with KR (pooled: *r* = −0.873, *P* < 0.001; SW *r* = −0.697, HM *r* = −0.948, HCP *r* = −0.969, HMCP *r* = − 0.942, all *P* < 0.05). Spectral gap (**B, E**) was positively correlated with MC within each network type (SW *r* = −0.988, HM *r* = −0.750, HCP *r* = 0.885, HMCP *r* = 0.856, all *P* < 0.01). The pooled correlation, however, was weak and non-significant (*r* = − 0.154, *P* = 0.317), a Simpson’s paradox in which between-type variation reverses the apparent direction of the within-type relationship when network types are aggregated. Non-normality (**C, F**) was positively correlated with both MC (pooled: *r* = 0.699, *P* < 0.001; per type: SW *r* = 0.875, HM *r* = 0.853, HCP *r* = 0.792, HMCP *r* = 0.832, all *P* < 0.01) and KR (pooled: *r* = 0.474, *P* < 0.001; HM *r* = 0.838, HCP *r* = 0.758, HMCP *r* = 0.859, all *P* < 0.01). Pooled random networks are not shown, as their inclusion would obscure topology-dependent patterns. Random networks showed strong correlations across all spectral–performance pairs (all |*r*| > 0.76, *P* < 0.001), confirming that the spectral mechanisms identified in structured networks generalize to unstructured counterparts.

hanges in kernel rank followed more complex patterns not explained by spectral radius alone. The reciprocity-driven decline in KR was robust to the choice of SVD threshold used in its computation, confirming that it reflects a genuine structural effect rather than a methodological artefact (Kernel rank computation; Supplementary Figures S12–S14). Across conditions, the spectral gap narrowed consistently with increasing reciprocity (Figure 2; Supplementary Figures S1 and S2), paralleling the decline in kernel rank, a relationship confirmed by formal correlation (*r* = −0.483, *P* < 0.001; Figure 5, panel **E**). We derived a theoretical result for linear reservoirs showing that under random Gaussian inputs, kernel rank is bounded by the dimension of a Krylov subspace, which expands as the spectral gap narrows (Supplementary Theorem 1). This result is consistent with the cross-architecture pattern: at a given reciprocity level, network types with a smaller spectral gap tend to exhibit higher kernel rank. Within each architecture, however, increasing reciprocity simultaneously narrows the spectral gap and reduces non-normality, and both effects jointly suppress KR.

Non-normality reflects the transient amplification of inputs arising from interactions between non-orthogonal eigenmodes. High non-normality enables reservoirs to represent inputs along many directions simultaneously, supporting richer dynamics and longer memory. As reciprocity increased and connectivity became more symmetric, eigenmodes became more orthogonal, trajectories aligned with dominant modes, and the representational space compressed. The combined reduction in spectral gap and non-normality explains the decline in kernel rank at high reciprocity, consistent with previous findings that non-normal networks sustain richer dynamics than normal ones with identical spectra (Hennequin et al., 2012; Ganguli et al., 2008).

Together, these results reveal a fundamental trade-off: reciprocity strengthens local feedback but restructures spectral properties in ways that suppress memory and representational richness. The MC decline reflects two contributing mechanisms, gain reduction from aggressive normalization and eigenspectral restructuring, while the KR decline is driven primarily by spectral gap narrowing and reduced non-normality. Since weight normalization is widely used in artificial recurrent networks, these reciprocity-related constraints may generalize well beyond reservoir computing.

### 2.8 Comparative advantages of network topologies shift with reciprocity, sparsity, and weight distribution

We next examined how reciprocity interacts with network topology, sparsity, and weight distribution. In binary networks and networks with narrow weight distributions, hierarchical modular (HM) architectures performed best at low reciprocity, significantly outperforming random networks in both memory capacity and kernel rank (*P* < 0.001; Figures 3, 4). With increasing reciprocity, however, HM performance converged toward random, reflecting the loss of long-range inter-modular links. In contrast, small-world (SW) networks gained a relative advantage under high reciprocity, particularly for memory capacity. Beyond link reciprocity = 0.4, SW structured networks not only diverged progressively from their randomized counterparts (Cohen’s *d* growing from 0.46 at reciprocity = 0.4 to 1.18 at reciprocity = 1.0, *P* < 0.001), but also performed comparably to or better than other structured architectures, suggesting that small-world topology confers a selective resilience to the computational costs of high reciprocity. Hierarchical modular core-periphery (HMCP) networks tracked random, while hierarchical core-periphery (HCP) consistently underperformed (Figures 3, 4; Supplementary Figures S3, S5 and S6). These results reveal a functional split: HM networks excel at low reciprocity, whereas SW networks are more resilient at high reciprocity.

In higher connectivity regimes, modular architectures (HM and HMCP) maintained advantages across reciprocity levels; memory capacity and kernel rank both peaked at intermediate values (Figure S7), reflecting pruning that balanced intra- and inter-modular communication and supported diverse internal dynamics.

Weight distribution further modulated these architectural trends. Broad log-normal weights enhanced baseline performance but steepened reciprocity-driven decline, particularly in lower connectivity networks. Under these conditions, architectural differences narrowed: modular and SW benefits diminished, HMCP resembled random, and HCP gained the most from weight heterogeneity, performing on par with other architectures (Figure 4; Supplementary Figures S5 and S6, lower panels).

Overall, these results demonstrate that the computational consequences of reciprocity depend jointly on network topology, sparsity, and weight distribution. Modular networks excel for low reciprocity, SW networks display selective resilience at high reciprocity, and weight heterogeneity reduces topological differences while exacerbating reciprocity-driven decline.

### 2.9 Reciprocity effects confirmed in empirical directed connectomes

Current noninvasive methods cannot reconstruct the human connectome as a directed network, and even large-scale comparative datasets provide only symmetric connectivity estimates (Suarez et al., 2022). To test whether the reciprocity effects observed in synthetic networks generalize to biology, we analyzed the only available directed connectomes of non-human primates: long-distance pathways of the macaque brain (Young, 1993; Sporns and Zwi, 2004; Hilgetag et al., 2000), the macaque visual cortex (Markov et al., 2014), and the cortical connectivity of the common marmoset (Majka et al., 2016).

These datasets differ in size, density, and reciprocity. The macaque long-distance connectome (47 nodes, density = 0.23, link reciprocity = 0.76) is binary and relatively sparse, with a mean out-degree of ∼10. For completeness, we also evaluated the earlier 71-node version (density = 0.15, reciprocity = 0.83; Supplementary Figure S15). By contrast, the macaque visual cortex (29 nodes) and marmoset cortex (55 nodes) are weighted, densely connected (densities = 0.66 and 0.63), and show intermediate strength reciprocity (0.55 and 0.53, respectively). Thus, the macaque connectome allowed us to test link reciprocity, while the visual and marmoset connectomes enabled analysis of strength reciprocity.

For each network, reciprocity was systematically varied and compared against 100 degree-preserving rewired null models, with spectral radii optimized for memory performance (Suárez et al., 2024). As shown in Figure 6, both empirical and null networks exhibited declining memory capacity and kernel rank with increasing reciprocity. The empirical macaque long-distance connectome consistently outperformed nulls across reciprocity values, and at its original reciprocity (0.76) achieved higher memory capacity than a rewired network with lower reciprocity (∼ 0.35) (*P* < 0.001, Cohen’s *d* = 0.89; Figure 6), likely reflecting its sparse, modular structure (see also Figure 3, panel **B**).

**Figure 6.**
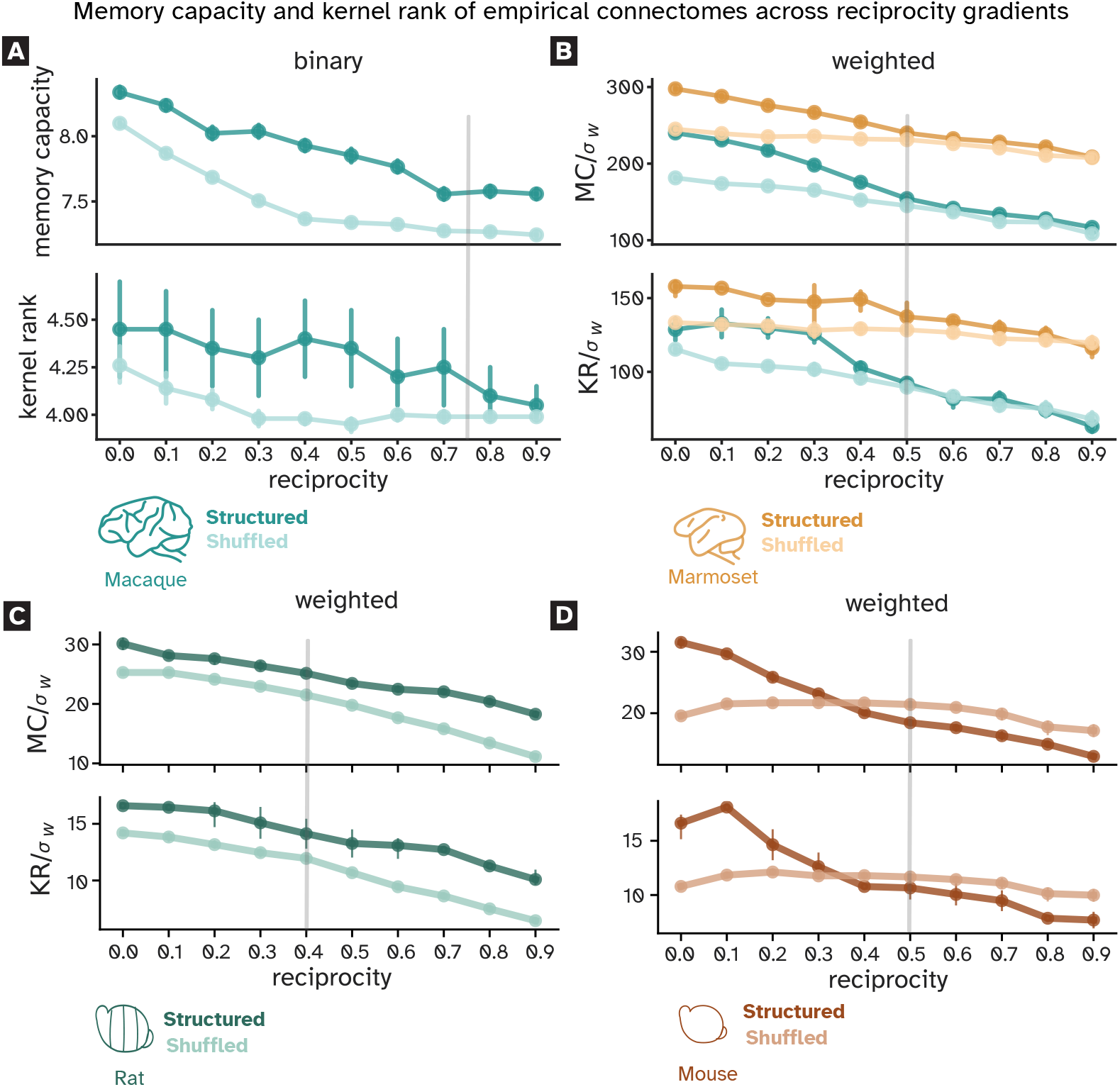
Memory capacity and kernel rank of empirical connectomes across reciprocity gradients. Panel **A** shows memory capacity and kernel rank as functions of link reciprocity for the binary macaque long-distance pathways connectome (47 nodes) and its degree-preserving rewired null models. Both metrics decline with increasing reciprocity, with the empirical connectome outperforming null models across all reciprocity values. At the original reciprocity (0.76), the reservoir achieved MC = 7.6 ± 0.2, versus 7.4 ± 0.2 for the rewired null at lower reciprocity (∼ 0.35; *P* < 0.001, Cohen’s *d* = 0.89). Panels **B, C**, and **D** show normalized memory capacity and kernel rank for the weighted macaque visual cortex, marmoset, rat, and mouse cortical connectomes and their null models. Normalization by the standard deviation of weights (σ_*w*_) accounts for weight redistribution effects accompanying increased strength reciprocity. Gray lines indicate empirical reciprocity values (macaque visual cortex and marmoset: *r* = 0.5; rat: *r* = 0.4; mouse: *r* = 0.5). Across all four weighted connectomes, both metrics declined significantly in structured networks. In macaque, marmoset, and rat, structured networks consistently outperformed their shuffled counterparts. In the mouse, the structured network outperformed shuffled at low reciprocity but the shuffled network exceeded it beyond reciprocity = 0.4. For the macaque visual cortex, MC/_σ*w*_ declined with slope = −120.4 (*R*^2^ = 0.938, *P* < 0.001) and KR/_σ*w*_ with slope = −65.7 (*R*^2^ = 0.862, *P* < 0.001). For the marmoset, MC/_σ*w*_ declined in the structured connectome (slope = −81.5, *R*^2^ = 0.919, *P* < 0.001) but not in the shuffled network (*R*^2^ = 0.183, *P* = 0.217); KR/_σ*w*_ followed a similar pattern (structured: slope = −41.3, *R*^2^ = 0.821, *P* < 0.001; shuffled: *R*^2^ = 0.067, *P* = 0.471). For the rat, both metrics declined near-linearly (MC/_σ*w*_: slope = −12.2, *R*^2^ = 0.988; KR/_σ*w*_: slope = −7.2, *R*^2^ = 0.968; all *P* < 0.001). For the mouse, structured declines were steep (MC/_σ*w*_: slope = −20.4, *R*^2^ = 0.961; KR/_σ*w*_: slope = −11.3, *R*^2^ = 0.910; all *P* < 0.001) while the shuffled network showed no significant trend (MC/_σ*w*_: *R*^2^ = 0.415, *P* = 0.045; KR/_σ*w*_: *R*^2^ = 0.397, *P* = 0.051).

In weighted connectomes, performance was lower at native reciprocity (∼ 0.5) than in rewired models with reduced reciprocity (≤0.15). The effect was pronounced in macaque (9.6 ± 0.3 vs. 11.6 ± 0.8) but minimal in marmoset (10.2 ± 0.3 vs. 10.3 ± 0.4), consistent with the marmoset’s larger overall weight sum and narrower distribution of weights. To control for redistribution effects introduced by reciprocity manipulation in weighted networks (Hadaeghi et al., 2026), memory capacity and kernel rank were normalized by weight variance. After normalization, both metrics declined with increasing reciprocity in empirical and null networks (Figure 6, panel **B**); non-normalized values in Supplementary Figure S15).

While our results indicate that lower strength reciprocity enhances computational performance in connectome-based reservoir computing models, the reciprocity values observed in macaque and marmoset connectomes are unlikely to be optimized solely for such performance. They likely reflect a trade-off between computational efficiency and other biological constraints. Our results suggest that an empirical reciprocity of ∼ 0.5 may mark a tipping point, beyond which empirical and rewired models perform similarly. One possible reason why reciprocity stabilizes around this intermediate level could be the laminar disentanglement of reciprocal projections: forward and feedback connections are distributed across different cortical layers, allowing substantial bidirectionality in the network as a whole while reducing the destabilizing effects of direct symmetric loops.

To probe cross-species generalization, we also analyzed directed weighted connectomes of rat (Bota et al., 2015) and mouse cortex (Rubinov et al., 2015). The rat connectome comprised 73 areas with a density of 0.37, strength reciprocity of 0.45, and link reciprocity of 0.66. The mouse connectome comprised 112 areas with a density of 0.53, strength reciprocity of 0.52, and link reciprocity of 0.67 (Figure 6, panels **C** and **D**). In the rat connectome, both normalized MC and KR declined monotonically with increasing reciprocity. Structured connectomes consistently outperformed their shuffled counterparts across all reciprocity values. The mouse connectome showed a different pattern, more similar to that observed in the marmoset connectome (Figure 6, panel **B**). Structured networks outperformed shuffled ones only at reciprocity values below approximately 0.4; above this threshold the advantage diminished. This is consistent with the native reciprocity of the mouse connectome (∼ 0.52) lying near a tipping point, as also noted for the marmoset connectome. These cross-species results suggest that the computational cost of reciprocity is not specific to the primate lineage, while also highlighting that the structural advantage of empirical connectomes depends on their native reciprocity level.

### 2.10 Reciprocity-driven performance decline generalizes across tasks, species, and network scales

To assess the robustness of our findings, we extended the analysis along three dimensions: task diversity, cross-species validation, and network scale.

To test whether the reciprocity effect generalizes beyond linear temporal recall, we evaluated reservoir performance on three additional tasks (see NARMA task evaluation and Context-dependent decision-making task): NARMA-5 and NARMA-10, which require nonlinear integration of past inputs over multiple timescales, and a context-dependent decision-making task (Mante et al., 2013), which requires selective integration across simultaneous input channels. For these experiments, we used random networks rather than bio-inspired architectures. This was a deliberate choice: by removing topology as a variable, we isolate the effect of reciprocity from the confounding performance advantages that structured architectures confer at low reciprocity (as documented in Sections 2.4–2.5). The cost of this design is that it does not characterize how the interaction between topology and reciprocity, a central finding of the main paper, extends to these additional tasks. Examining whether hierarchical modular or small-world architectures show differential resilience across nonlinear tasks remains an important direction for future work. Unless otherwise stated, all experiments in this section used networks with N = 400 nodes and *k* = 6. For the decision-making task, we additionally report results at *k* = 10 (Supplementary Figure S18). The interaction between link reciprocity and connectivity level across memory tasks is examined through a sparsity sweep with *k* = 5, 10, 20, 50 reported below.

Under random weight assignment, MC, NARMA-5, and NARMA-10 all declined significantly across both weight distributions, with steeper slopes and wider variance under broader weights (Supplementary Figures S16 and S S17). The decision-making task behaved differently. For random weights at *k* = 6, balanced accuracy dropped only modestly across the full reciprocity range (0.651 ± 0.018 to 0.629 ± 0.058), and the wide variance at high reciprocity indicates limited sensitivity to structural changes in the absence of weight symmetry enforcement. Under enforced strength reciprocity, however, the decline was larger and more consistent across all tasks. For the decision-making task at *k* = 6, balanced accuracy dropped from 0.652 ± 0.018 to 0.546 ± 0.030 and macro F1 from 0.647 ± 0.021 to 0.515 ± 0.043 (Supplementary Figure S18 and Supplementary Table S3). This pattern mirrors the results for MC and the NARMA tasks. A weak effect under random weights becomes pronounced under enforced strength reciprocity, consistent with strength reciprocity being the more potent of the two reciprocity forms for degrading computational capacity. Under binary connectivity, MC declined clearly and consistently (slope = −2.78, *R*^2^ = 0.996, *P* < 0.001). NARMA-5 showed a shallow but statistically significant decline (slope = −0.007, *P* < 0.001), while NARMA-10 exhibited a steeper nonlinear profile, declining sharply at low reciprocity and flattening beyond *r* ≈ 0.5 (slope = −0.050, *R*^2^ = 0.853, *P* < 0.001). The decision-making task showed a negligible change under binary connectivity, likely because this task places greater demands on representational capacity, leaving less room to detect reciprocity-driven degradation at this network size and connectivity level.

We further examined how connectivity level modulates the reciprocity effect using binary networks with *N* = 400 nodes across *k* = 5, 10, 20, and 50 (Supplementary Figure S19 and Supplementary Table S4). Sparser networks showed superior MC and stronger reciprocity-driven decline across all three tasks. For MC, increasing reciprocity produced a significant negative linear trend at all out-degrees (all *P* < 0.001), but the effect attenuated progressively with density: the slope decreased from −4.03 at *k* = 5 to −0.11 at *k* = 50, with the latter representing a difference of less than one unit across the full reciprocity range. For NARMA-5 and NARMA-10, the reciprocity effect was restricted to the sparsest regime: only *k* = 5 showed a significant negative trend (both *P* < 0.001), while *k* = 10 and *k* = 50 showed no significant change across reciprocity levels (all *P* > 0.06). At *k* = 20, a small but statistically significant positive trend emerged in both NARMA tasks (NARMA-5: *P* < 0.001; NARMA-10: *P* = 0.010), suggesting that in moderately dense networks, higher reciprocity marginally improves nonlinear task performance. These results confirm that the reciprocity effect on working memory is strongest in the sparse regime, consistent with our main findings, and that this regime is most relevant to both biological neural circuits and resource-constrained artificial systems.

Cross-species generalization to rat and mouse cortical connectomes is reported in Section 2.9.

Finally, extending synthetic networks to *N* = 512 and *N* = 1024 nodes confirmed that reciprocity-driven decline persists at larger scales, with qualitatively identical trends to those observed at *N* = 64–256 (Supplementary Tables S1 and S2), arguing against a small-network artifact.

## 3 Discussion

### Summary

Our study provides a systematic and mechanistic account of how reciprocity, in both its structural (link) and weighted (strength) forms, constrains the computational capacity of recurrent neural networks. This work offers a functional explanation for a long-standing observation in cortical neuroanatomy: the systematic asymmetry of reciprocal connections, whereby feedforward projections are consistently stronger than their feedback counterparts. Across a wide range of network architectures, including biologically inspired, empirical, and random topologies, we show that increasing reciprocity consistently impairs memory capacity and representational richness as measured by kernel rank, two core indicators of a network’s ability to support complex, high-dimensional dynamics.

Such reciprocity-induced functional impairments are not limited to binary wiring. They persist, and in some cases worsen, with the introduction of biologically plausible heterogeneity in connection weights using log-normal distributions. While greater weight variability can enhance absolute memory performance (Damicelli et al., 2022), it does not buffer the system against the negative effects of reciprocity. On the contrary, in networks with longer-tailed weight distributions, the computational cost of reciprocal topology becomes more severe. Strikingly, when both link and strength reciprocity are enforced, mimicking conditions with abundant bidirectional motifs, we observe an exponential decline in memory and kernel rank across all architectures. These results suggest a fundamental computational pitfall that biological systems may have evolved to avoid.

The functional decline associated with strong direct loops can be traced to the spectral structure of the network. We show that the spectral radius, driven by a dominant positive eigenvalue, increases with reciprocity, pushing the network toward instability. To preserve the echo-state property and maintain fair comparisons, we applied spectral normalization to all networks, tuning them to operate near a common dynamical regime with a spectral radius close to 1.0. However, higher reciprocity required more aggressive weight suppression to reach this regime, resulting in a marked reduction in total connection strength. This reduction along with eigenvalue restructure, likely underlies the observed loss in memory capacity. Crucially, even under matched spectral conditions, high-reciprocity networks consistently underperformed, revealing that reciprocity imposes a structural limitation on the usable parameter space of recurrent systems. The relationship between cycle structure and reservoir computational capacity has been examined analytically in prior work (Peng et al., 2024; Aceituno et al., 2020), where eigenvalue distribution arguments provide a foundation for understanding how loop topology shapes memory and representational richness. Our results complement and extend such findings to the weighted reciprocity regime.

These results are consistent with theoretical work on recurrent excitatory-inhibitory networks showing that reciprocal motifs shift leading eigenvalues and destabilize dynamics (Shao and Ostojic, 2023; Dahmen et al., 2020). We note, however, that those excitatory-inhibitory network models operate under Dale’s Law constraints with sign-segregated excitatory and inhibitory populations, whereas our models use non-negative weights only. The sign-constrained spectral structure of biological EI circuits, therefore, lies outside our model’s scope. Nevertheless, the spectral shift observed here, in which reciprocity elevates the spectral radius and narrows the spectral gap, is mechanistically analogous to the feedback-driven instability described in those frameworks.

The decline in representational diversity, measured by kernel rank, followed more complex and architecture-specific patterns. Across conditions, the spectral gap tended to narrow with increasing reciprocity, correlating with reduced kernel rank. Although not strictly linear, this decline suggests that reciprocity compresses transient network dynamics. While kernel rank is known to depend on spectral radius and input scaling (Legenstein and Maass, 2007), its link to topology has been less explored. We derived a theoretical result for linear reservoirs showing that the decline in kernel rank with reciprocity may arise from the combined influence of a narrowing spectral gap and changes in eigenvalue non-normality, pointing to an important direction for future theoretical work. This complements studies using Lyapunov spectra, which link coupling and input to attractor dimensionality and entropy, and low-rank models showing how rapidly decaying singular values compress effective dimensionality (Engelken et al., 2023; Thibeault et al., 2024).

Beyond spectral and dimensional effects, our results also reveal that network architecture interacts with reciprocity in distinct ways, with differences emerging across sparsity regimes and weight distributions. In binary and narrowly weighted networks, modular and hierarchical topologies outperformed random and small-world ones in the lower connectivity regime, particularly at low reciprocity. In the higher connectivity regime, they retained an advantage across reciprocity levels, often showing a clear peak in memory and representational capacity at intermediate reciprocity. Bio-inspired structures appear to place reciprocal edges where they are most useful, such as within modules or along hierarchical pathways, which helps to offset some of the performance costs of reciprocity in lower connectivity regimes. With broad weight distributions, structural advantages are no longer evident, and the negative impact of reciprocity follows a linear decline in the higher connectivity regime, but accelerates into an exponential decay in the lower connectivity regime. By considering both regimes, we capture the conditions most relevant to different classes of networks. In particular, the higher connectivity regime is more relevant for biological neural circuits, while the lower connectivity regime is more relevant for artificial neural networks. This framework highlights how density, reciprocity, and network topology jointly define computational capacity.

### Relevance to neuroscience

The present findings, validated for the available directed empirical connectomes spanning four mammalian species (macaque, marmoset, rat, and mouse), offer a computational rationale for the no-strong-loop principle observed in cortical circuits. Specifically, in sparse and metabolically constrained biological systems, minimizing reciprocity may serve to maintain functional stability while preserving dynamical richness. These findings align with recent experimental work showing that biological neural cultures exhibit highly sample-efficient adaptation, a property thought to arise from their sparse yet rich connectivity patterns (Khajehnejad et al., 2025, 2023).

Our results also connect to theoretical work on how network motifs relate to different kinds of memory in cortical circuits. Brunel (Brunel, 2016) showed that networks optimized for long-term associative memory, via stable attractor states, tend to over-represent strong reciprocal connections. By contrast, networks supporting the dynamical maintenance of sequential activity patterns show no such bias. This distinction highlights that reciprocity plays different roles across different memory regimes. Classical attractor models, including Hopfield-type networks (Hopfield, 1982; Krotov and Hopfield, 2016), rely on symmetric connectivity to stabilize fixed-point states and support robust long-term storage. Recent evidence that enriched experience increases synaptic reciprocity in non-sensory association cortex and hippocampus (Saxena et al., 2025) is consistent with this picture, as these regions support pattern completion and associative retrieval. Here, we focus on a different regime: rather than modelling working memory as a cognitive process, we use memory capacity as a task-agnostic measure of a network’s ability to retain recent sequential inputs, a computational resource that underlies, but is not equivalent to, the full complexity of working memory in biological systems, which encompasses not only maintenance but also active transformation and manipulation of retained content (Nyberg and Eriksson, 2016). In this regime, strong reciprocity reduces non-normal dynamics and degrades memory capacity and representational diversity, revealing a fundamental trade-off between stable associative storage and flexible transient maintenance and processing. These perspectives are complementary rather than contradictory. Recurrent circuits may generate high-dimensional internal dynamics while task-relevant activity occupies lower-dimensional manifolds (Jazayeri and Ostojic, 2021), and different brain regions may exploit this division. In particular, hippocampal circuits implicated in long-term memory favor stronger reciprocal motifs (Saxena et al., 2025; Jensen et al., 2024), while distributed cortical circuits supporting working memory favor weaker direct reciprocity to preserve dynamical richness. Moreover, our results show that this constraint operates independently of other structural motifs, because reciprocity-induced degradation appears even in random architectures. However, certain topologies, such as hierarchical core-periphery networks, exhibited greater resilience when weight variability was high, suggesting that evolutionary pressures may have favored architectures that buffer against reciprocity’s trade-offs.

A crucial distinction emerging from the present findings is that, while recurrence is generally essential for neural computation (Goldman-Rakic, 1995; Wang, 2001; Miller et al., 2018; Saxena and Cunningham, 2019), not all forms of recurrence are equally advantageous. Our results indicate that direct reciprocity, in the form of strong and symmetric bidirectional connections, tends to degrade memory and representational richness. By contrast, recurrence mediated through loops or chains of connections likely supports stable yet flexible

dynamics (Crick and Koch, 1998; Hennequin et al., 2012; Shao and Ostojic, 2023). This pattern resonates with the observed cortical organization, where complementary forward and feedback projections between areas are common but characteristically segregated across different laminar circuits, creating extended recurrent loops, rather than direct bidirectional motifs. Such an arrangement preserves the computational advantages of recurrence while avoiding the destabilizing effects of symmetry. The area-intrinsic segregation of feedforward and feedback pathways may also be the reason for the observed relatively high level of connectional symmetry in the analyzed empirical data sets, which is likely compensated for by laminar disentanglement. It remains an important challenge to test whether similar segregation mechanisms apply across spatial scales, particularly in local microcircuits examined by dense connectomics (Seeman et al., 2018; Motta et al., 2019; Peng et al., 2024). Extending the analysis to invertebrate and non-mammalian vertebrate nervous systems is a natural future direction. The Drosophila connectome (Scheffer et al., 2020; Lin et al., 2024) and larval zebrafish whole-brain reconstructions (Hildebrand et al., 2017; Légaré et al., 2025) are compelling candidates. However, such analyses would test whether the computational consequences of asymmetric connectivity generalize across radically different circuit architectures — not whether the no-strong-loops hypothesis holds in those systems. That hypothesis is mechanistically grounded in the laminar organisation of cortical projections, a specificity established in primates and rodents but not in non-mammalian systems. Recurrent excitatory connections are sparse in mouse and human cortical microcircuits (Seeman et al., 2018), and synaptic connectivity in human layer 2-3 cortex tends toward acyclic organisation (Peng et al., 2024). Whether these patterns reflect an analogous computational constraint or a different organisational principle remains open.

An important future challenge is to examine how reciprocity gradients are organized along cortical axes and whether they help balance stability and representational diversity. Cortical systems already show large-scale gradients in connectivity, excitability, and timescales that support hierarchical processing (Goulas et al., 2018; Fulcher et al., 2019; Goulas et al., 2021). Our results suggest that spatial variations in reciprocity may provide a complementary mechanism, tuning recurrent dynamics to the demands of different regions. This question can now be addressed with dense connectomic datasets across species and scales, from larval zebrafish whole-brain reconstructions (Légaré et al., 2025; Hildebrand et al., 2017) and the Drosophila connectome (Scheffer et al., 2020; Winding et al., 2023; Lin et al., 2024) to mammalian cortical microcircuits (Seeman et al., 2018; Motta et al., 2019; Shapson-Coe et al., 2024). Such resources enable reciprocity to be probed both within local circuits and across inter-areal pathways, and related to principles such as laminar structure, core–periphery topology, and excitation–inhibition balance. These integrative analyses promise deeper insight into how the cortex achieves both reliable stability and flexible, high-dimensional computation.

Beyond structural connectomes, asymmetric reciprocity can be characterized in any directed network representation of brain organization. Whether similar computational consequences emerge when reciprocity is varied in directed functional or effective connectivity estimates (Mill et al., 2017; Reid et al., 2019) is an open question.

### RELEVANCE to NeuroAI and further fields

These findings also hold practical implications for artificial recurrent systems. Many models in AI and neuromorphic computing use randomly initialized recurrent layers, often with minimal structural constraints. As interest grows in sparsely connected, brain-inspired designs, our findings suggest that reciprocity should be treated as an independent control parameter, not just an incidental property of random connectivity. This view resonates with recent work in artificial architectures, such as Transformers, where directional constraints in attention patterns have been shown to influence learning and representation dynamics (Rathor et al., 2025; Saponati et al., 2025). Managing reciprocity may help optimize stability, memory, and representational diversity in artificial systems, particularly those operating in resource-constrained environments or aiming to emulate biological efficiency.

#### Limitations and conclusions

Several limitations should be noted alongside the deliberate methodological choices that motivated them. The reservoir computing framework isolates the contribution of network structure to computational capacity by keeping recurrent weights fixed and training only the readout layer. This separation is a feature rather than a limitation: it allows us to attribute differences in memory capacity and representational diversity directly to reciprocity and topology, without conflating structural effects with those of learning dynamics. Models in which learning occurs within the recurrent layer, through biologically plausible plasticity rules such as reward-modulated Hebbian learning (Miconi, 2017) or gradient-based optimization (Murray, 2019), constitute a natural and important next step. Our results establish the structural baseline from which such models can depart: if reciprocity imposes spectral constraints on the computational resources available to a fixed network, those same constraints are likely to shape the trajectory and outcome of within-network learning. The present framework is nonetheless a simplification. It does not generate behavior, and differs from task-optimized recurrent networks trained to perform cognitive tasks (Yang et al., 2019), as well as from activity flow models that simulate neural information transfer over empirically estimated functional connectivity (Ito et al., 2022; Mill et al., 2022). These are complementary rather than competing approaches: where task-optimized and activity flow models ask how known connectivity supports specific cognitive functions, our approach asks how structural motifs shape the computational resources available irrespective of task. Beyond the learning assumption, several further scope limitations apply. Theoretical analyses were derived mainly for linear reservoirs; although simulations suggest similar effects in nonlinear settings, a full nonlinear theory is still lacking. The empirical connectomes analyzed, though spanning rodents and non-human primates, are incomplete, biased toward certain regions and species, and may not capture the full diversity of reciprocity. Extrapolating the present conclusions to very large trained recurrent networks, where weight distributions and spectral properties are shaped by gradient-based optimization across millions of parameters, also requires caution, as the dynamical regime may differ substantially from the sparse, fixed-weight reservoirs studied here. Our models also assumed fixed connectivity and input statistics, whereas biological circuits are plastic and modulated by neuromodulators. Finally, while the implications for artificial intelligence remain to be tested directly in state-of-the-art architectures, Figure 7 summarizes the key computational consequences of reciprocity and their potential translational relevance for both biological and artificial recurrent networks.

**Figure 7.**
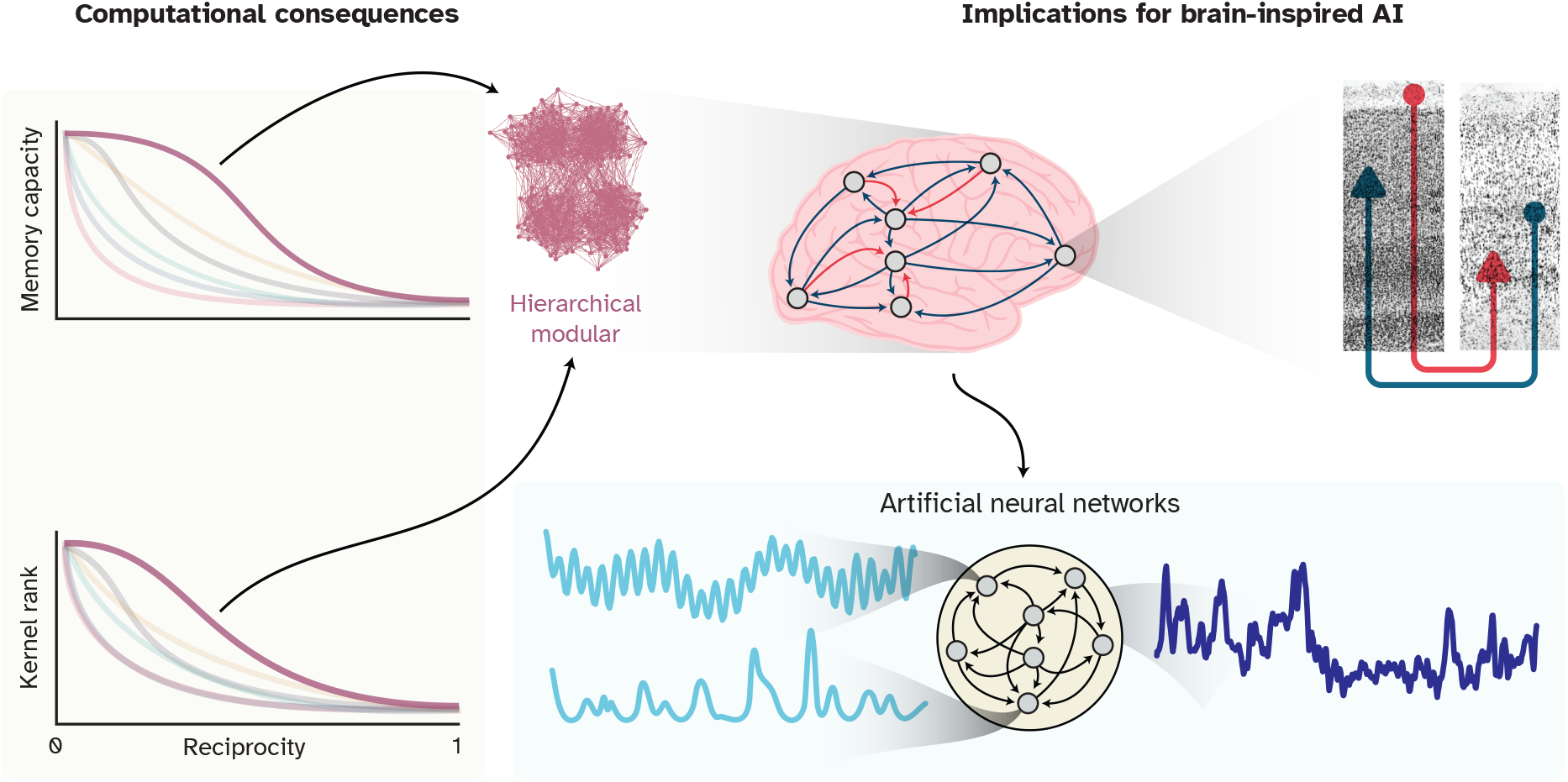
Reciprocity as a structural principle in neural and artificial networks. Reciprocity is a fundamental topological property that systematically constrains memory capacity and representational richness in recurrent networks, with its impact shaped by architecture, connection density, and weight distribution. In neuroscience, gradients of reciprocity, together with the laminar segregation of forward and feedback pathways, may tune local dynamics while mitigating destabilizing symmetry, a prediction now testable with emerging detailed, multi-scale connectomic datasets. For artificial and neuromorphic systems, reciprocity may be treated as a tunable design feature: low reciprocity enhances stability and memory, while task-specific levels may unlock richer architectures. More broadly, managing reciprocity as a structural parameter provides a route toward next-generation neural network designs that balance stability with computational richness, rather than inheriting the inefficiencies of random connectivity.

In summary, we demonstrate that reciprocity is a key feature of neural network organization. It imposes intrinsic constraints on memory and dynamic diversity in recurrent networks, even under optimal spectral tuning. By linking this constraint to underlying spectral mechanisms, we provide a unifying explanation for the suppression of reciprocity in brain circuits and offer actionable design principles for the next generation of efficient, brain-inspired artificial networks. These findings bridge neuroscience, network theory, and machine learning, showing how structural motifs shape functional capacity across both natural and artificial systems.

## 4 Materials and Methods

### 4.1 Generating synthetic network architectures

We constructed four classes of directed binary networks: small-world (SW), hierarchical modular (HM), hierarchical core-periphery (HCP), and hierarchical modular core-periphery (HMCP), alongside matched random networks serving as null models. All networks were generated at five sizes (*N* = 64, 128, 256, 512 and 1024 nodes) under two sparsity regimes, corresponding to mean out-degrees of *k* = 6 (lower connectivity) and *k* = 10 (higher connectivity).

#### Small-world networks

We used the Watts-Strogatz algorithm (Watts and Strogatz, 1998), implemented via the NetworkX library, to generate small-world topologies. The algorithm begins with a ring of *N* nodes, each connected to its *k* nearest neighbors by undirected edges. Each edge is then rewired with probability *P* to a randomly selected target node, avoiding self-loops and duplicate edges. To ensure small-world properties, we selected intermediate values of *P*: specifically, *P* = 0.4 for lower connectivity networks (*k* = 6) and *P* = 0.5 for higher connectivity networks (*k* = 10). As this algorithm produces undirected graphs, we used our network reciprocity control (NRC) algorithms, detailed in the Reciprocity adjustment algorithms section and (Hadaeghi et al., 2026), to introduce directionality and adjust reciprocity.

#### Hierarchical modular networks

Hierarchical modular networks were constructed using the recursive algorithm developed by Sporns (Sporns and Zwi, 2004), implemented through the bctpy library (Rubinov and Sporns, 2010). Each network was represented by a directed, binary adjacency matrix of size *N* = 2^*l*^, with *l* ∈ {6, 7, 8, 9, 10} corresponding to 64, 128, 256, 512, and 1024 nodes. The network is initialized with fully connected elementary modules of 2^*s*^ nodes. These modules are recursively merged, with inter-module connection density decreasing by a factor of *E*^−*s*^ at hierarchical level *s*. We tuned *E* and *s* to match the desired link density and maximize memory capacity. For lower connectivity (*k* = 6) networks, we used *s* = 4, *E* = 4.0; for higher connectivity (*k* = 10) networks, *s* = 2, *E* = 3.5. Reciprocity was subsequently adjusted using the NRC algorithm.

#### Hierarchical core-periphery and modular core-periphery networks

We extended the hierarchical modular model to incorporate core-periphery structure by embedding centrality gradients into the construction process. Hierarchical core-periphery (HCP) topology was initialized with a star-like seed motif, biasing early nodes to form a densely interconnected core with stronger projections to peripheral nodes. During recursive construction, we enforced differential connectivity templates: core nodes exhibited high mutual connectivity and dominant out-degree, while peripheral nodes had lower intra-group connectivity and weaker projections overall.

To formalize this topology, we computed a hierarchical connection cost matrix *H* ∈ ℝ ^*N*×*N*^, where *H*_*ij*_ reflects the hierarchical level and topological distance at which nodes *i* and *j* become connected. For HMCP networks, we applied a 10% attenuation to intra-cluster connectivity in peripheral modules to maintain modular structure. The connection probability was defined as:

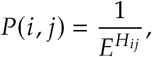

where *E* is a tunable decay parameter.

Parameter values for *E* and *s* were optimized to match the intended link density and enhance memory performance. lower connectivity networks (*k* = 6) used *s* = 2, *E* = 4.0 for HCP and *s* = 2, *E* = 4.0 for HMCP networks; higher connectivity networks (*k* = 10) used *s* = 4.0, *E* = 3.25. In higher connectivity networks, we set *s* = 4, *E* = 1.61 for HCP and *s* = 3, *E* = 3.83 for HMCP networks. Final networks were adjusted using the NRC algorithm for specified reciprocity levels.

#### Synthetic random baselines

As computational baselines for the synthetic architectures, we generated random directed binary networks *de novo* rather than by rewiring the structured graphs. For each network of size *N*, every node was assigned exactly *k* outgoing connections (target mean out-degree), with targets sampled uniformly at random without replacement from all other nodes, excluding self-connections and duplicate edges. This procedure yields fixed out-degree (directed *k*-out) random graphs that match the structured networks in size and overall density, while leaving in-degree distributions, reciprocity, and higher-order topology unconstrained.

We intentionally adopt this fully random construction, rather than degree-preserving rewiring, because standard reservoir computing practice relies on fixed randomly connected reservoirs which, when appropriately scaled, provide strong generic computational performance and therefore constitute a natural baseline against which structured architectures can be compared (LukoševiIčius, 2012). After generating binary connectivity, weights were assigned using the same allocation schemes as for the structured networks to ensure comparability across architectures (See Assigning connection weights). For each structured network instance, five matched null networks were independently generated, ensuring robust estimation of the performance advantage of structured over random connectivity.

#### Empirical connectome randomization

For empirical connectomes, where only a single observed topology is available, we instead constructed topology-randomized counterparts by rewiring the original graph using the randmio_dir_connected function from the bctpy library. This procedure preserves global density and strength statistics while randomizing connection partners and maintaining network connectedness. These randomized networks serve as topology-controlled nulls, allowing us to assess how perturbations of the wiring of a fixed biological connectome influence computational performance. Comparisons of empirical connectomes to fully random reservoir baselines were examined separately in prior work Damicelli et al. (2022).

### 4.2 Assigning connection weights

To generate weighted networks from binary adjacency matrices, we assigned positive connection strengths drawn from a log-normal distribution. This choice is supported by empirical evidence indicating that large-scale cortical connectivity, especially area-to-area projection strengths, follows heavy-tailed, positively skewed distributions well described by log-normal statistics (Markov et al., 2014; Theodoni et al., 2022; Piazza et al., 2025).

Weights were assigned only to existing (non-zero) entries in the binary adjacency matrix, thereby preserving the structural sparsity and topology of the original network. For each network instance, we calibrated the log-normal distribution to match a specified target mean *µ*_*t*_ and standard deviation σ_*t*_ of connection strengths. To do so, we computed the corresponding parameters of the underlying normal distribution in log-space as:

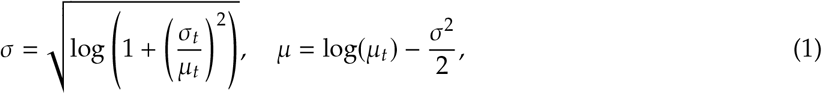

where *μ* and σ are the mean and standard deviation of the associated normal distribution in logarithmic space. Using these parameters, weights were independently sampled for each existing connection. Random seeds were fixed to ensure reproducibility across trials.

This approach allowed us to generate weighted networks that retained the binary topological structure while incorporating biologically plausible heterogeneity in connection strengths. Spectral normalization was subsequently applied as described in the Reservoir computing framework section to ensure dynamical stability across network instances.

### 4.3 Reciprocity adjustment algorithms

To manipulate reciprocity, we employed the network reciprocity control (NRC) framework developed in our previous work (Hadaeghi et al., 2026). The same algorithms are used across synthetic and empirical networks, but in different operational modes reflecting the degree of experimental control available. For synthetic networks, reciprocity is treated as a design parameter: we first generate binary topologies using the chosen generative model, then adjust link reciprocity via NRC, subsequently assign weights, and finally adjust strength reciprocity using the weighted NRC procedure. This two-stage process enables independent control of both structural (link-level) and strength (weight-level) reciprocity. In contrast, empirical connectomes are available only as fixed binary or weighted networks; consequently, NRC is applied post hoc to adjust link or strength reciprocity while preserving the observed topology and global weight statistics. Thus, in synthetic networks, NRC serves as a generative control mechanism, whereas in empirical networks, it functions as a controlled adjustment of an existing biological structure. Below, we summarize the binary and weighted formulations used in both settings.

#### 4.3.1 Binary reciprocity

In binary networks, link reciprocity quantifies the proportion of bidirectional edge pairs relative to the total number of directed edges. For a binary adjacency matrix, *A*, it is defined as:

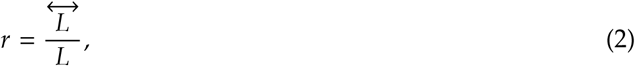

where *L* denotes the total number of directed edges, and 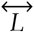 is the number of reciprocal edge pairs, i.e., instances where both (*i* → *j*) and (*j* → *i*) exist.

To achieve a target reciprocity level *r*_*t*_, we use a discrete rewiring procedure that maintains total edge count. If the current reciprocity *r*_*c*_ exceeds the target, reciprocal pairs are selectively broken and reassigned to unconnected node pairs. Conversely, if *r*_*c*_ < *r*_*t*_, unidirectional edges are selectively converted into reciprocal pairs. For each new reciprocal link added, an existing unidirectional edge elsewhere in the network is removed, ensuring that the total number of directed edges, and thus the network density, remains unchanged. Throughout this process, self-loops are prohibited, and the directed, binary structure is preserved. Rewiring continues iteratively until the network’s reciprocity reaches the target value within a predefined tolerance.

#### 4.3.2 Weighted reciprocity

In weighted (directed) networks, let *W* = [*w*_*ij*_] denote the adjacency (weight) matrix; reciprocity quantifies the symmetry of connection strengths between pairs of nodes. Following the framework of Squartini et al. (Squartini et al., 2013), the symmetric (reciprocated) weight between nodes *i* and *j* is:

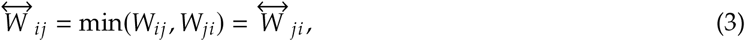

and the asymmetric (non-reciprocated) component is:

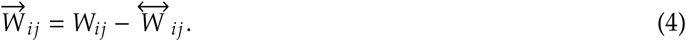

The overall strength reciprocity of a weighted network is then:

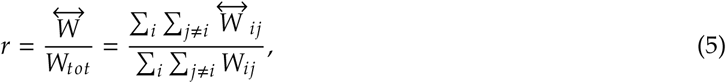

Where 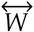 denotes the sum of all symmetric weight components and *W*_*tot*_ is the sum of all connection weights in the network.

To steer a network toward a target strength reciprocity *r*_*t*_, we apply a controlled rescaling procedure. When decreasing reciprocity, symmetric weights are scaled down, and the difference is proportionally redistributed to the asymmetric components, preserving total weight. When increasing reciprocity, symmetric components are gradually amplified while adjusting asymmetric weights accordingly, ensuring all values remain non-negative. The algorithm terminates once the weighted reciprocity reaches the target value within a specified tolerance.

Together, these procedures allow for independent and precise adjustment of both structural and strength reciprocity across diverse network architectures. This control enables systematic exploration of how reciprocal motifs influence dynamics and computation, both in biologically inspired and abstract artificial networks. Full algorithmic details and complexity analysis are provided in (Hadaeghi et al., 2026).

### 4.4 Spectral property analysis

To understand how graded patterns of reciprocity shape the global dynamical characteristics of directed networks, we analyzed key spectral features of the adjacency matrices. These features, including spectral radius, spectral gap, and Henrici’s index of non-normality (Asllani et al., 2018), provide insight into how structural properties affect signal propagation, amplification, and stability, especially in recurrent systems.

#### Spectral radius

The spectral radius of a matrix, *W*, is defined as the largest absolute value among its eigenvalues:

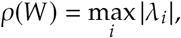

where λ_*i*_ are the eigenvalues of *W*. Recurrent neural networks, including both fixed-weight reservoir systems and trainable vanilla RNNs, operate as complex dynamical systems. In such systems, the spectral radius is a critical control parameter that governs the stability and richness of temporal dynamics. When weights are fixed, as in reservoir computing, the spectral radius directly determines whether the system satisfies the echo-state property (Jaeger and Haas, 2004) and remains stable over time. In trainable RNNs, where weights are optimized through backpropagation through time, the spectral radius of the recurrent weight matrix plays a key role in gradient stability: values significantly greater than one can cause gradients to explode, while values significantly less than one can lead to vanishing gradients, both of which hinder effective learning (Pascanu et al., 2013).

#### Spectral gap

The spectral gap captures the separation between the dominant eigenvalues and is defined as:

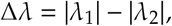

assuming that eigenvalues are ordered such that |λ_1_| ≥ |λ_2_| . The spectral gap has been linked to the rate of information decay and the richness of transient dynamics in recurrent networks.

#### Henrici’s index of non-normality

To quantify the degree of non-normality in the network’s adjacency matrix, we computed Henrici’s departure from normality. The Frobenius norm of a matrix, *W*, is given by:

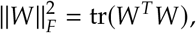

and the Henrici deviation is:

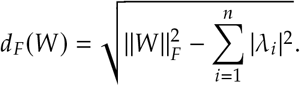

This value is zero for normal matrices and increases with non-normal structure. To ensure comparability across networks of different sizes and densities, we used the normalized Henrici index:

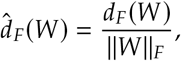

as proposed in (Asllani et al., 2018). This index ranges from 0 (perfectly normal matrix) to 1.0 (maximally non-normal), providing a scale-invariant measure of the matrix’s asymmetric and non-orthogonal eigenvalue structure.

### 4.5 Reservoir computing framework

Reservoir computing (RC) is a biologically inspired paradigm for processing temporal signals using recurrent neural networks (RNNs) (Jaeger and Haas, 2004; Maass et al., 2002). It consists of three components: an input layer, a fixed recurrent reservoir, and a readout layer. The core idea is that the reservoir’s internal weights are fixed and untrained, enabling the system to project inputs into a high-dimensional, dynamic state space with minimal computational cost.

This fixed-weight architecture is particularly suited for investigating how network topology influences computation, as it decouples learning from structural adaptation. Only the readout layer is trained, typically via linear regression, transforming the learning task into a convex and efficient optimization problem. As such, RC offers a tractable framework for probing how structural properties such as modularity, sparsity, and reciprocity affect the computational capacity of recurrent systems (Rodriguez et al., 2019; Damicelli et al., 2022; Achterberg et al., 2023; Fakhar et al., 2022).

In our experiments, we adopted a leaky integrator (LI) neuron model to govern reservoir dynamics (Jaeger et al., 2007). The LI formulation introduces a leak rate parameter α ∈ [0, 1], which controls the temporal inertia of reservoir units. Higher α values correspond to faster responses to incoming signals, while lower values introduce slower dynamics. The state update equation is:

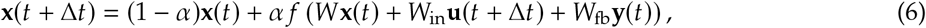

Where 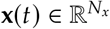is the reservoir state at time, *t*, 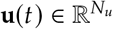 is the input, 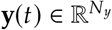 is the output, and *f* is the element-wise nonlinearity (here, *t an h*). The matrices *W, W*_in_, and *W*_fb_ denote the recurrent, input, and feedback weight matrices, respectively. The discrete time step Δ*t* determines the update interval and is typically chosen based on the timescale of the input dynamics. In this study, we set Δ*t* = 1. Moreover, we used α = 1, which led to improved memory capacity across tested configurations.

The reservoir output at time *t* is given by:

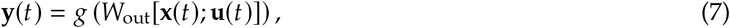

where [·;·] denotes vector concatenation, *g* is a nonlinear output activation, and *W*_out_ is the trainable readout matrix. In this study, the output activation function, *g*, was set to a linear function. It allows the readout layer to directly combine reservoir states and inputs without additional nonlinearity.

To determine *W*_out_, we used ordinary least squares (OLS) regression with Tikhonov regularization:

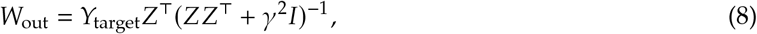

Wher 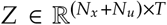 is the matrix of concatenated reservoir and input states (i.e., [**x** (*t*) ; **u** (*t*)]) across *T* time steps, 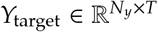 contains the temporal target output signals, *I* is the identity matrix, and *γ* ≥ 0 is a regularization parameter. Regularization improves numerical stability and guards against overfitting, especially in noisy or high-dimensional settings. We used *γ* = 10^−5^ in all simulations.

All simulations were implemented using the open-source echoes package (Damicelli et al., 2022).

### 4.6 Memory capacity estimate

To quantify the short-term memory capacity of each network, we followed a standard delayed memory task. In each trial, the network received a one-dimensional input signal, *u*(*t*), generated as a sequence of independent, identically distributed samples from a standard normal distribution 𝒩(0, 1).

The network was trained to reconstruct delayed versions of this input using separate output units, each targeting a different delay *n*. Specifically, the desired output for unit *n* was defined as:

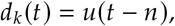

Where *n* = 1, 2, …, 1.4*N*_*x*_ (Farkaš et al., 2016), and *N*_*x*_ is the number of reservoir nodes.

Following training, we evaluated predictions on a held-out test sequence using the coefficient of determination, *R*^2^, as implemented in scikit-learn and adopted by the conn2res toolbox(Suárez et al., 2024). The memory capacity (MC) was then computed as:

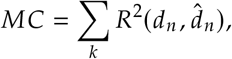

where *d*_*n*_ and 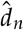 denote actual and predicted delayed inputs, respectively. This measure reflects how well the reservoir retains and separates past inputs across multiple timescales, with higher MC indicating superior memory performance. We note that the classical definition of memory capacity uses the squared Pearson correlation coefficient (Farkaš et al., 2016); however, *R*^2^ has been increasingly adopted in the neuroscience community for reservoir computing applications (e.g., conn2res and (Milisav et al., 2025)), and yields comparable results when predictions are accurate.

Each evaluation used a sequence of *T* = 3,200 time-steps, split into 70% training and 30% test sets in temporal order. Input weights were drawn from Uniform (™1.0, 1.0) and scaled by a factor of 0.05. Small independent Gaussian noise (σ = 10^−4^) was added to reservoir states during training to improve generalization.

### 4.7 Kernel rank computation

To quantify the representational capacity of each reservoir network, we computed the kernel rank (KR), following procedures adapted from prior work in reservoir computing (Legenstein and Maass, 2007; Dale et al., 2021). KR reflects the network’s ability to project diverse inputs into distinct internal states, a property closely linked to the separation principle in reservoir computing (Maass et al., 2002).

For each network, we constructed a matrix

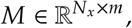

where each column vector **x**_*ui*_ represents the reservoir state evoked by a distinct input stream *u*_*i*_, and *N*_*x*_ denotes the number of reservoir units.

We then applied singular value decomposition (SVD) to *M*, yielding singular values σ_1_ ≥ σ_2_ ≥ · · · σ_min (*Nx,m*)_ . The kernel rank was defined as the number of singular values exceeding a relative threshold:

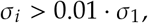

where σ_1_ is the largest singular value. This thresholding approach reduces the influence of noise or minor components and captures only the dominant axes of variation in the reservoir state space.

To ensure sufficient coverage of the input space and stable KR estimates, we set the number of input samples equal to the reservoir size, i.e., *m* = *N*_*x*_.

### 4.8 NARMA task evaluation

To assess whether the effect of reciprocity on computational performance extends beyond linear temporal recall, we evaluated reservoir networks on two nonlinear autoregressive moving average tasks: NARMA-5 and NARMA-10 (Rodan and Tino, 2010). These tasks require the network to reproduce a target sequence whose current value depends nonlinearly on both past inputs and past outputs over a horizon of *n* = 5 or *n* = 10 timesteps, respectively. They are canonical benchmarks for probing nonlinear memory in recurrent systems because they cannot be solved by any linear readout operating on instantaneous inputs alone.

The target sequence *y*(*t*) was generated according to the recurrence:

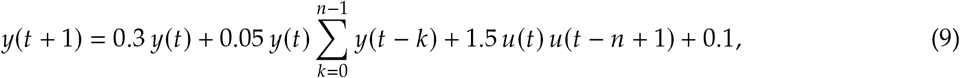

where *u* ∼ (*t*) Uniform (0, 0.5) is the driving input and *n* is the memory length (5 or 10). A warm-up period of 200 time-steps was discarded before recording the sequence to avoid initialization artifacts. Sequences that diverged to non-finite values were regenerated with a different random seed, with up to 50 retries per trial.

Each evaluation used a sequence of *T* = 1,000 time-steps, split into 70% training and 30% test sets in temporal order. The reservoir received a one-dimensional input *u* (*t*) and was trained to predict *y* (*t* + 1) one step ahead. A transient of 300 time-steps was discarded from the training states to eliminate initial reservoir dynamics, and a further 100 time-steps were discarded from the test states before computing performance metrics. Reservoir states and inputs were concatenated to form the feature vector presented to the readout.

Experiments were conducted on networks with a random architecture of *N* = 400 nodes with a mean out-degree *k* = 6. Weight assignment and reciprocity adjustment followed the same procedures as described in Sections 4.2 and 4.3, respectively. For each of the 11 reciprocity values (*r* ∈ {0.0, 0.1, …, 1.0}), 50 independently generated networks were evaluated, each with 5 independent input weight matrix seeds, and results were averaged across runs. The spectral radius was set to 0.95 for both tasks, input weights were drawn from Uniform(−1.0, 1.0) and scaled by a factor of 0.3. Small independent Gaussian noise (σ = 10^−4^) was added to reservoir states during training to improve generalization.

Performance was quantified using the Pearson correlation coefficient between predicted and target outputs on the held-out test segment, computed after the test washout period. This metric was chosen for consistency with the memory capacity measure reported in the main analyses. Values close to 1.0 indicate accurate reproduction of the target sequence; values near 0 indicate failure to capture the nonlinear temporal structure of the task.

### 4.9 Context-dependent decision-making task

To probe whether reciprocity effects generalize to tasks requiring nonlinear integration of multi-channel inputs and context-dependent classification, we evaluated reservoirs on the context-dependent decision-making task introduced by Mante et al. (Mante et al., 2013), as implemented in the NeuroGym and conn2res toolboxes.

In this task, the network receives inputs across seven simultaneous channels at each time-step (Δ*t* = 100 ms): a fixation signal, a context or rule cue, and two pairs of stimulus channels representing competing sensory evidence in two modalities. Each trial proceeds through a fixation period (300 ms), a stimulus period (750 ms), a variable delay, and a decision period (100 ms). The network must selectively integrate evidence from the relevant sensory modality as indicated by the context cue and report a three-class decision (fixate, choose left, or choose right). The task, therefore, requires the simultaneous maintenance of context information, selective gating of sensory evidence, and nonlinear integration across input channels. These properties are absent from both the memory capacity and NARMA tasks.

Data were generated for 1,000 trials and concatenated into a single temporal sequence. The sequence was split 70/30 into training and test sets in temporal order. Reservoir states were collected using the leaky integrator dynamics described in Section 4.5, with input weights of shape (*N*× 7) drawn from Uniform (™1.0, 1.0) and scaled by a factor of 0.1. A transient of 15 time-steps was discarded from the training states. Reservoir states were concatenated with the corresponding input channels to form the feature vector at each time-step.

Because the task requires classification rather than regression, the readout was implemented as a ridge classifier trained on feature vectors from the decision period only, defined as time-steps at which the target label was either class 1 (choose left) or class 2 (choose right). Fixation-period time-steps (class 0) were excluded from both training and evaluation to focus the classifier on the decision-relevant portion of the trial. Performance was quantified using balanced accuracy on the held-out test set, which accounts for potential class imbalance between left and right choices. Macro-averaged F1 score was computed as a secondary metric.

Experiments used networks with random architecture and *N* = 400 nodes with mean out-degree *k* = 6 and *k* = 10. The spectral radius was optimized and set to 1.2, reflecting the finding that context-dependent tasks benefit from dynamics near or slightly above the edge of stability (Suárez et al., 2024).

### 4.10 Statistical analysis

To evaluate the computational performance of each reservoir network architecture, we computed the mean memory capacity (MC) at each level of reciprocity, averaging across 50 independently generated network instances and 5 training trials per instance. For each condition, we reported both the mean and standard deviation of the MC and KR values to capture central tendency and variability (Supplementary Table S1).

To assess whether structured networks outperformed their random counterparts, we conducted pairwise comparisons of MC and KR values using two-tailed independent-samples *t*-tests. Statistical significance was accompanied by effect size estimates based on Cohen’s *d*, providing a standardized measure of the magnitude of observed differences (Supplementary Table S1). For networks with *N* = 1024, Welch’s t-test was used in place of Student’s t-test to account for unequal variances arising from the reduced number of trials and null networks at this scale.

To quantify trends in performance decline with increasing reciprocity, we applied linear regression to estimate the slope of the decline in MC and KR across reciprocity values. The slope reflects the change in performance per unit reciprocity (scale 0 → 1), and its statistical significance was assessed via the *F*-test associated with the linear model. *R*^2^ quantifies goodness of fit. For the adjusted-broad weight regime (σ = 0.9), where performance decline was nonlinear, an exponential model (*y* = *a* · *e*^*b*·*x*^ + *c*) was fitted to mean MC as a function of reciprocity. The decay rate *b* and its statistical significance were estimated from the standard error of *b* via a *t*-test. Regression-based statistics are tabulated in Supplementary Tables S1 and S2.

For nonlinear memory tasks (NARMA-5 and NARMA-10), performance was quantified as Pearson correlation between network output and target, and summarized as mean and standard deviation across network instances (Supplementary Table S4). For the context-dependent classification task (Mante et al., 2013), performance was quantified using balanced accuracy and macro F1 score (Supplementary Table S3).

All statistical analyses were performed using standard scientific computing libraries in Python. A significance threshold of *P* < 0.05 was used for all hypothesis tests.

## Acknowledgments

We thank Gorka Zamora-López, Changsong Zhou, Petra Vértes, Marcus Kaiser, and Ben Fulcher for their valuable feedback and discussions, which helped improve the clarity and rigor of this work.

## Author contributions

F.H. and C.H. conceptualized and designed the study. F.H. developed the algorithms and conducted the formal analysis. F.H. and K.F. analyzed the data. K.F. created the graphs and visualizations. M. K. conducted the theoretical results. All authors contributed to organizing and writing the manuscript.

## Data and materials availability

The data supporting the findings of this study are publicly available. The connectome dataset can be accessed via Netneurotools athttps://netneurotools.readthedocs.io. Synthetic data and code used for implementing the algorithm and conducting network analysis in this study are available on GitHub. Specifically, the Network Reciprocity Control (NRC) algorithms can be found in the NRC repository, and the memory capacity and kernel rank computation codes are provided in the memory-capacity-and-kernel-rank repository.

## Supplementary materials

Supplementary theoretical results, Figures S1 to S19, and Tables S1 to S4.

## Supplementary Materials

## Supplementary theoretical results: spectral gap and kernel rank in linear reservoirs

### Problem Setup

We consider a linear reservoir computing system defined by the recurrent equation:

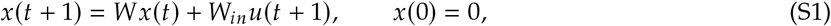

where:

- *x*(*t*) ∈ R^*N*^ is the hidden state at time *t*,
- *W* ∈ R^*N*×*N*^ is the recurrent weight matrix,
- *W*_*in*_ ∈ R^*N*×*d*^ maps input *u*(*t*) ∈ R^*d*^ into the reservoir.

Given a fixed evaluation time *t*^*∗*^ and *m* distinct input sequences 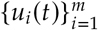 of length *T*, each producing a final hidden state *x*_*i*_(*t*^*∗*^), we define the reservoir state matrix:

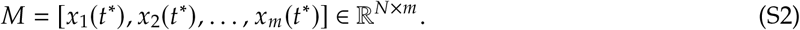

#### Definition 1

(Kernel Rank). *The* kernel rank *at time t*^*∗*^ *is defined as* rank (*M*), *representing the number of linearly independent reservoir responses to the set of inputs*.

The goal is to understand how the spectrum of *W* and its structure (e.g., symmetry, non-normality) influence this rank.

### Connection to Krylov Subspaces

Each hidden state can be expressed as:

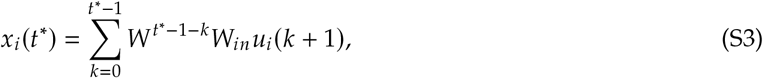

indicating that all *x*_*i*_ (*t*^*∗*^) lie in the span of vectors of the form *W* ^*k*^*W*_*in*_*u*, for 0 ≤*k* < *t*^*∗*^. This motivates the use of Krylov subspaces.

#### Definition 2

(Krylov Subspace). *Given W* ∈ R^*N*×*N*^ *and W*_*in*_ ∈ R^*N*×*d*^, *the Krylov subspace of order t*^*∗*^ *is:*

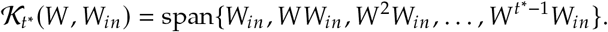

#### Lemma 1

(Krylov Rank Controls Kernel Rank). *Let r* = dim 𝒦_*t*_^*∗*^ (*W, W*_*in*_). *Then:*

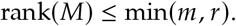

*Moreover, if the inputs u*_*i*_(*t*) ∼ 𝒩(0, σ *I*) *i.i.d. for all t, i, then:*

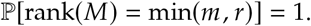

*Proof*. Each *x*_*i*_ *t*^*∗*^ is a linear combination of vectors in 𝒦_*t*_^*∗*^ (*W*), (*W*_*in*_), hence lies in this subspace. The rank of *M* is therefore bounded by *r*. For Gaussian input streams, the projected inputs Φ*u*_*i*_ (with Φ a matrix built from *W* ^*k*^*W*_*in*_) span the image of Φ with probability 1, achieving the maximum rank.

### Spectral Gap and Krylov Growth

Let *W* be diagonalizable: *W* = *Q*Λ*Q*^−1^, with eigenvalues {λ_*i*_} ordered by magnitude |λ_1_| ≥ |λ_2_| ≥ · · ·≥ |λ_*n*_| . Define the *spectral gap* as:

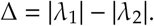

#### Theorem 1

(Spectral Gap Controls Kernel Rank). *Let W be diagonalizable and u*_*i*_(*t*) *be i.i.d. Gaussian inputs. Then, as the spectral gap* Δ → 0, *the expected kernel rank* E [(rank *M*)] *increases, provided W*_*in*_ *is full-rank and not aligned with a single eigenvector*.

*Sketch*. From above, rank *M* is upper-bounded by dim 𝒦_*t*_^*∗*^ (*W*), (*W*_*in*_) . The Krylov subspace consists of vectors *Q*Λ^*K*^ *Q*^−1^*W*_*in*_ for *k* = 0 to *t*^*∗*^™ 1. If Δ is large, repeated powers of Λ decay all but the dominant mode, leading to alignment with a single eigenvector direction. If Δ is small, multiple eigenmodes decay slowly, contributing diverse directions to the span, thereby increasing the rank. Hence, a smaller spectral gap leads to higher-dimensional Krylov subspaces, which implies a higher expected kernel rank.

### Role of Non-Normality and Symmetry

The above analysis assumes only eigenvalue magnitudes influence rank growth. However, the geometry of eigenvectors plays a critical role. In particular:

- If *W* is symmetric (normal), it is orthogonally diagonalizable, and each eigenvector evolves independently.
- If *W* is non-normal, its eigenvectors may be non-orthogonal, allowing for transient interactions and amplification that expand the Krylov subspace more rapidly.

Therefore, even with the same spectrum (including spectral gap), symmetric (high-reciprocity) matrices tend to produce lower kernel rank than asymmetric (low-reciprocity) matrices, due to reduced non-normality.

This theoretical framework explains our experimental observation that at low reciprocity, networks with a smaller spectral gap exhibit comparatively higher kernel ranks, reflecting richer internal representations. However, as reciprocity increases, the resulting rise in symmetry reduces non-normality, suppressing interactions between eigenmodes even as the spectral gap continues to narrow. This interaction leads to a decline in representational diversity despite slower decay dynamics. Together, these results highlight a fundamental trade-off: both spectral gap and non-normality jointly govern the expressivity of linear reservoirs, with symmetry imposing geometric constraints that can counteract the benefits of prolonged temporal integration.

### Implications

These results offer a theoretical foundation for the empirical link between spectral structure and kernel rank observed in the main text. Specifically: i) The spectral gap limits the number of persistent modes contributing to the hidden state. ii) Reciprocity constrains the directions in which those modes evolve. iii) Both factors must be considered to explain the nonlinear relationship between reciprocity and representational richness in recurrent networks.

**Figure S1.**
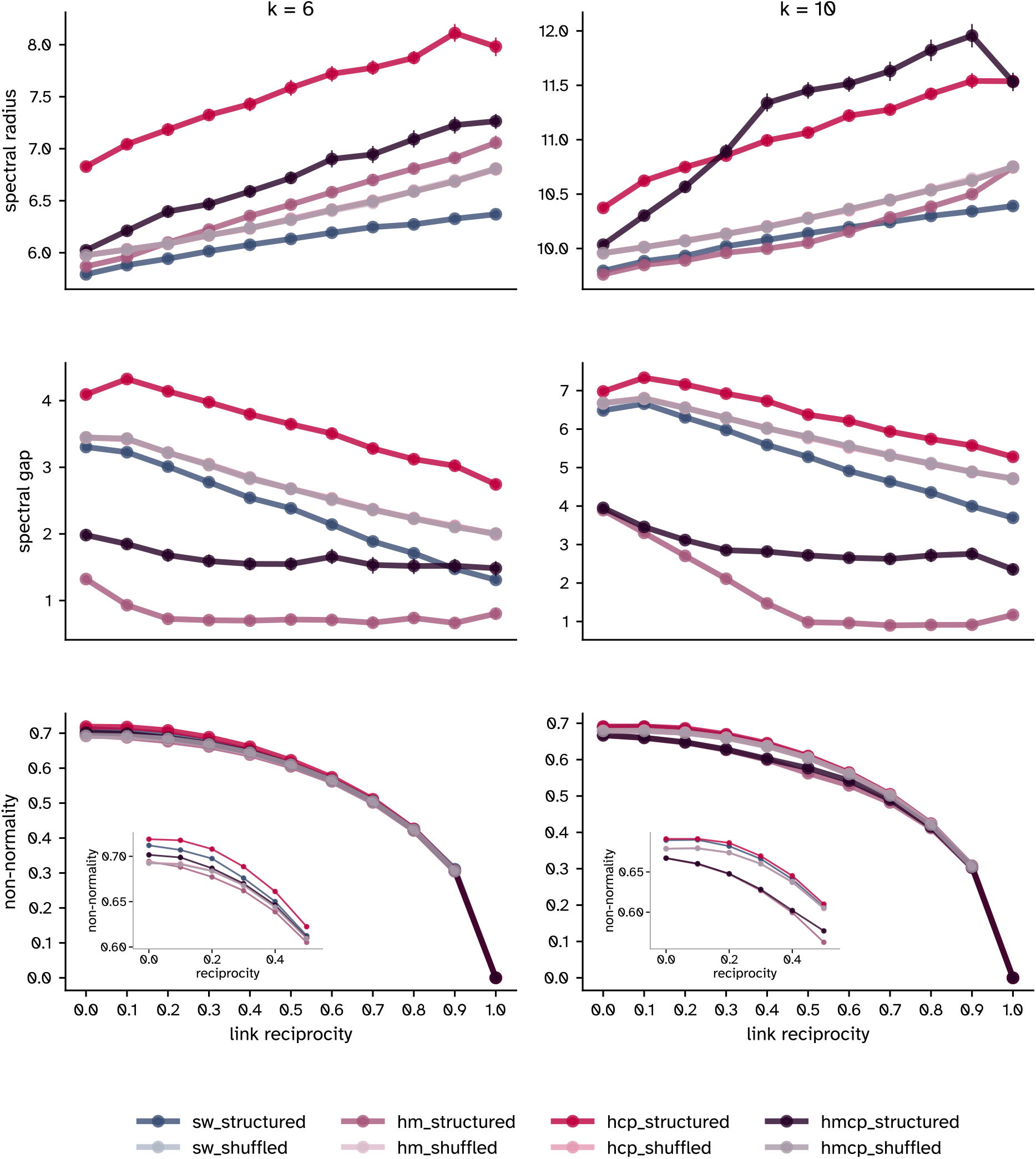
Spectral properties of synthetic networks across reciprocity gradients (128-node networks). Shown are the spectral radius, spectral gap, and Henrici’s index of non-normality for binary networks with 128 nodes under lower connectivity (mean out-degree *k* = 6, left) and higher connectivity (mean out-degree *k* = 10, right). Each point represents the mean across 50 instances; error bars denote standard deviation. Insets in the non-normality panels show an expanded view of the low reciprocity range (0–0.5), where architecture-dependent differences are most pronounced before convergence at higher reciprocity levels. Results are consistent with those observed for 64-node networks (Figure 2), confirming that increasing reciprocity systematically restructures the spectral properties of diverse network architectures across network sizes.

**Figure S2.**
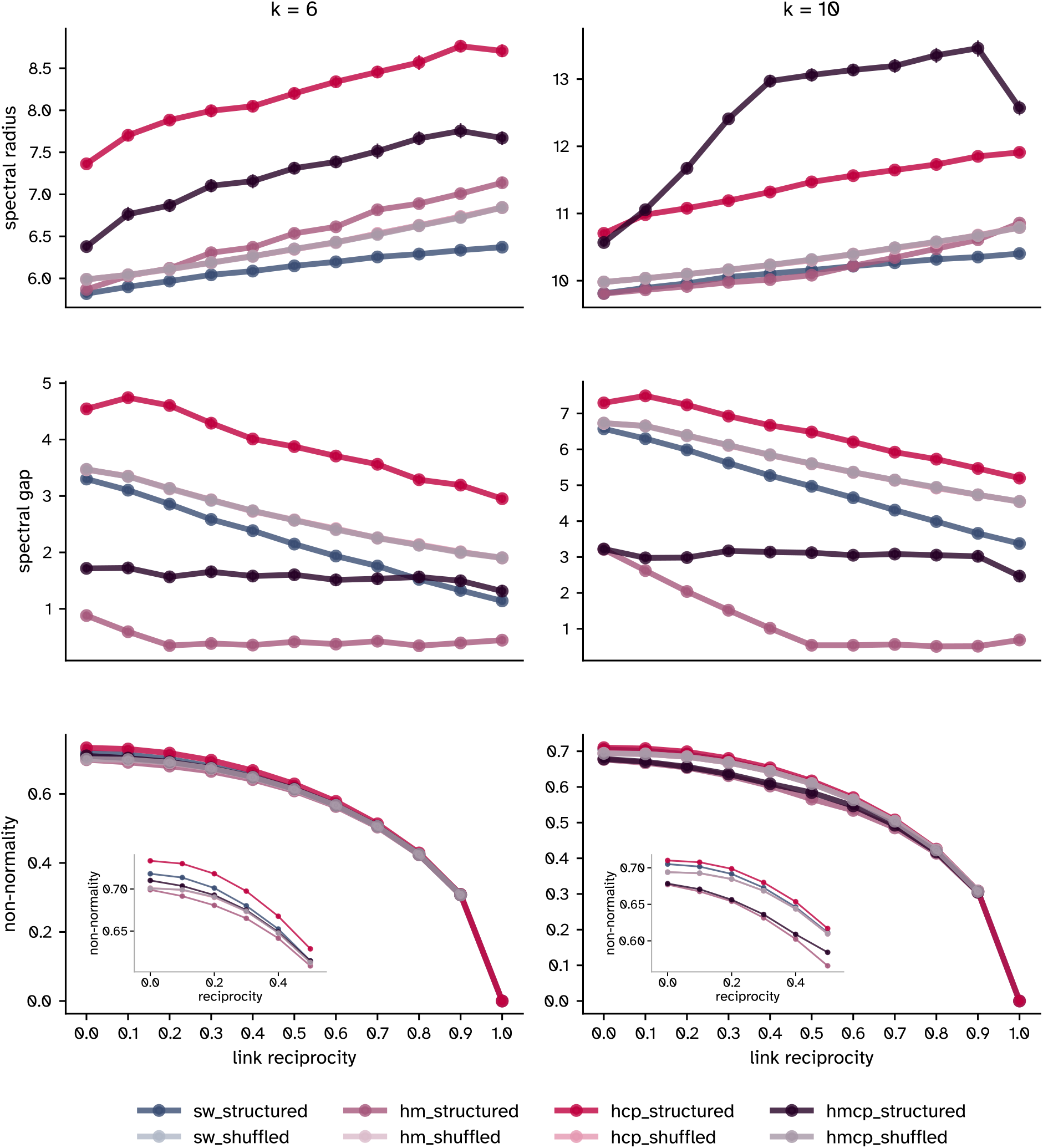
Spectral properties of synthetic networks across reciprocity gradients (256-node networks). Shown are the spectral radius, spectral gap, and Henrici’s index of non-normality for binary networks with 256 nodes under lower connectivity (mean out-degree *k* = 6, left) and higher connectivity (mean out-degree *k* = 10, right). Each point represents the mean across 50 instances; error bars denote standard deviation. Insets in the non-normality panels show an expanded view of the low reciprocity range (0–0.5), where architecture-dependent differences are most pronounced before convergence at higher reciprocity levels. Results are consistent with those observed for 64-node networks (Figure 2), confirming that increasing reciprocity systematically restructures the spectral properties of diverse network architectures across network sizes.

**Figure S3.**
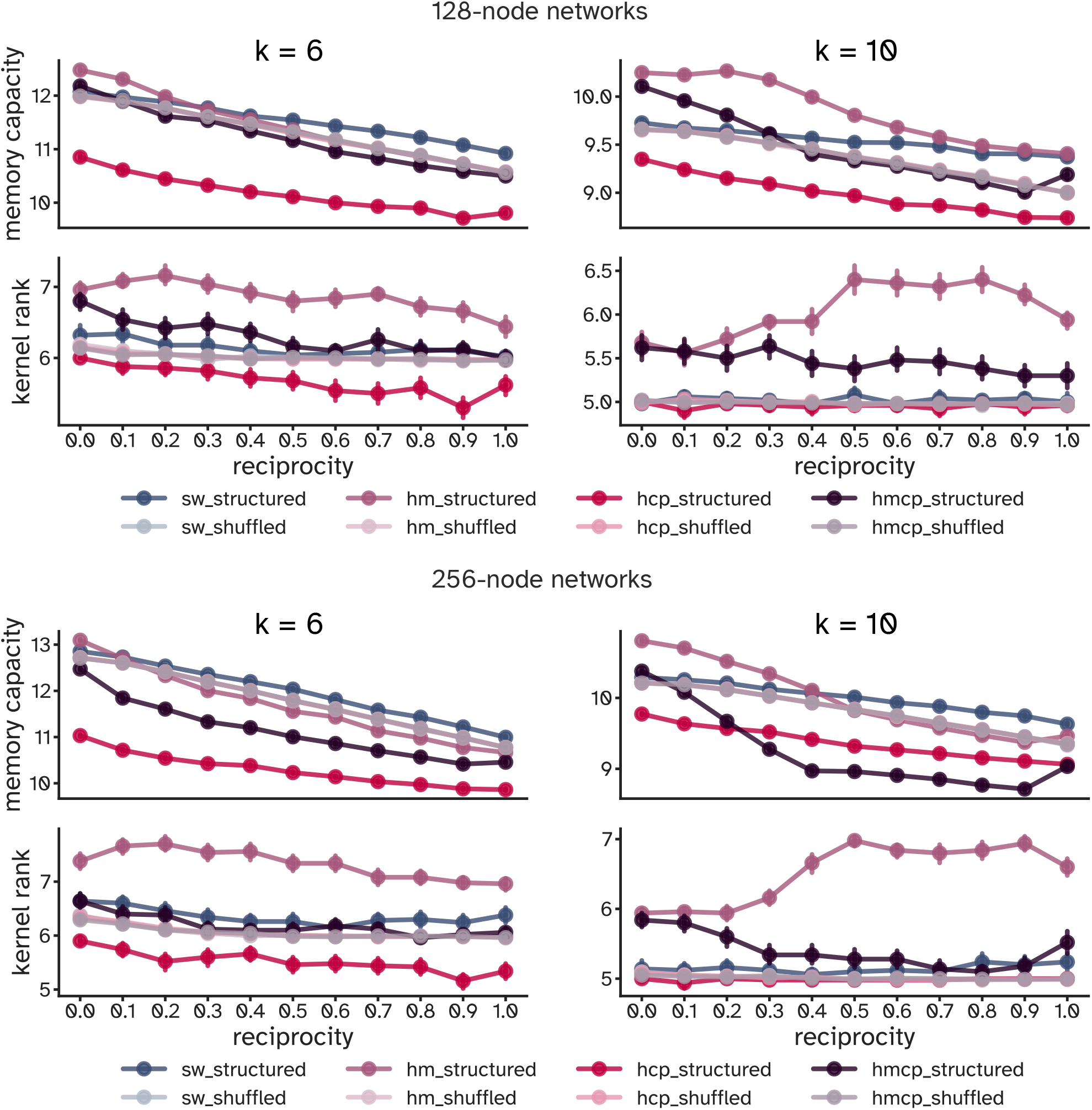
Link reciprocity reduces memory capacity and kernel rank in recurrent networks. Mean memory capacity (MC) and kernel rank are plotted as functions of link reciprocity for binary networks with 128 nodes (top panel) and 256 nodes (bottom panel), evaluated under both lower connectivity and higher connectivity regimes. Each point represents the average over 50 independently generated network realizations; error bars indicate standard deviation. While most architectures show a consistent decline in both MC and kernel rank with increasing reciprocity, kernel rank in hierarchical modular networks under the higher connectivity regime exhibits a non-monotonic trend, peaking at reciprocity = 0.5. Here, network generation parameters were held constant and matched those optimized for 64-node networks, allowing for cross-size comparisons under fixed topological constraints.

**Figure S4.**
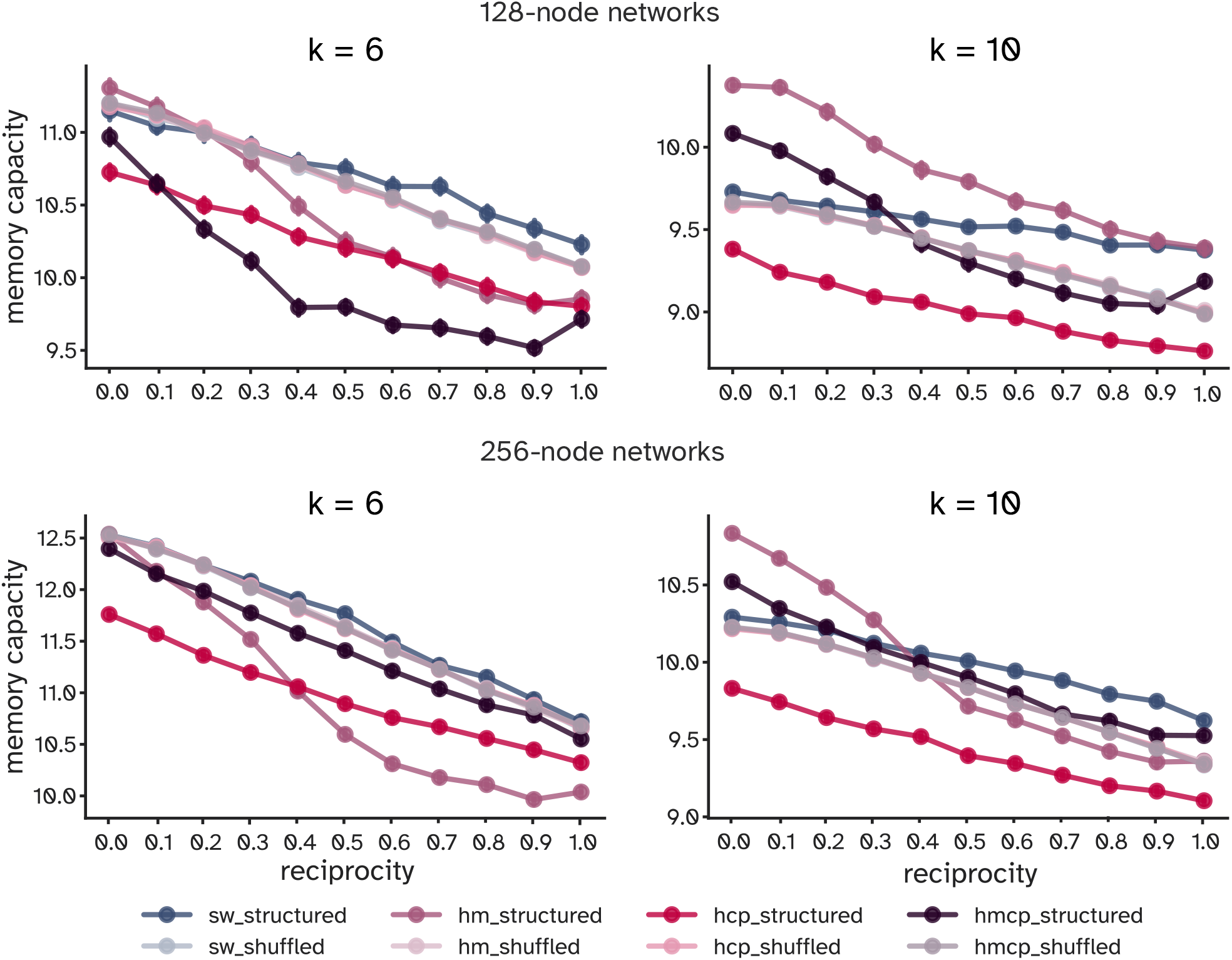
Optimized large-scale networks confirm reciprocity-driven computational decline. Memory capacity (MC) is shown as a function of link reciprocity for binary networks with 128 nodes (top panel) and 256 nodes (bottom panel), across lower connectivity and higher connectivity regimes. Each data point reflects the average over 50 independently generated networks; error bars denote standard deviation. In contrast to the previous figure, network generation parameters were re-optimized for each network size to maximize memory capacity. Although performance differences among architectures vary from those observed under fixed parameters, the decline in MC with increasing reciprocity remains consistent across all architectures. Parameter set for each network architecture with 128 nodes is as follows: small-world wiring probability = 0.4, hierarchical modular networks: *l* = 7, *E* = 3.72, *s* = 2.0, hierarchical core-periphery networks: *l* = 7, *E* = 1.52, *s* = 3.0, and hierarchical core-periphery networks: *l* = 7, *E* = 3.87, *s* = 2.0. Parameter set for each network architecture with 256 nodes is as follows: small-world wiring probability = 0.4, hierarchical modular networks: *l* = 8, *E* = 4.56, *s* = 2.0, hierarchical core-periphery networks: *l* = 8, *E* = 1.63, *s* = 3.0, and hierarchical core-periphery networks: *l* = 8, *E* = 1.87, *s* = 3.0.

**Figure S5.**
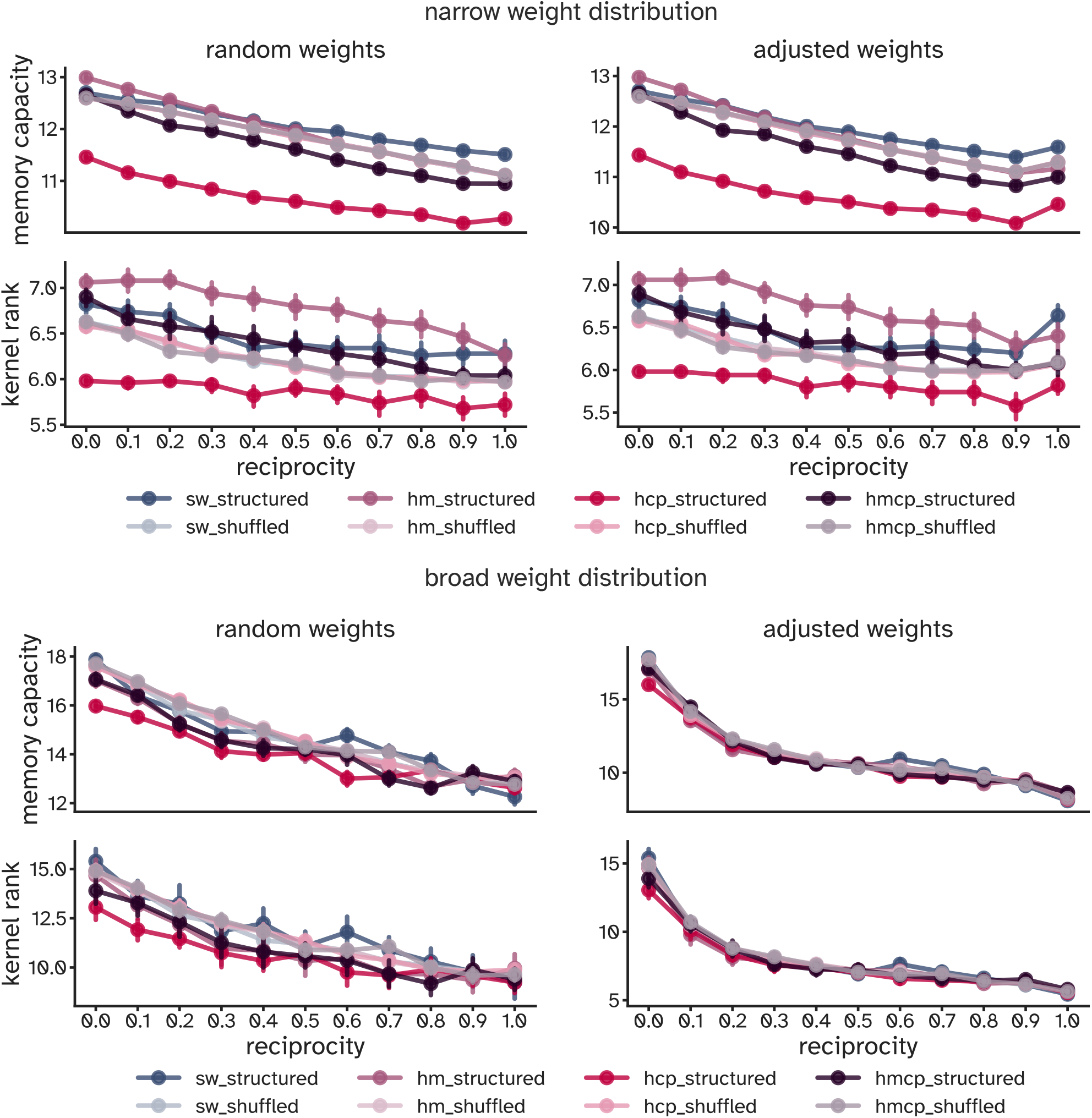
Reciprocity-driven declines in memory and kernel rank persist across weighted 128-node networks. Each panel compares performance under random (left) and adjusted (right) strength reciprocity, for weight distributions with standard deviations of 0.1 (top) and 0.9 (bottom), mean = 0.25. Memory capacity (MC) and kernel rank (KR) both decline as link reciprocity increases, especially under high weight variability. Patterns are consistent with those observed in 64-node networks. Error bars reflect standard deviation over 50 network realizations.

**Figure S6.**
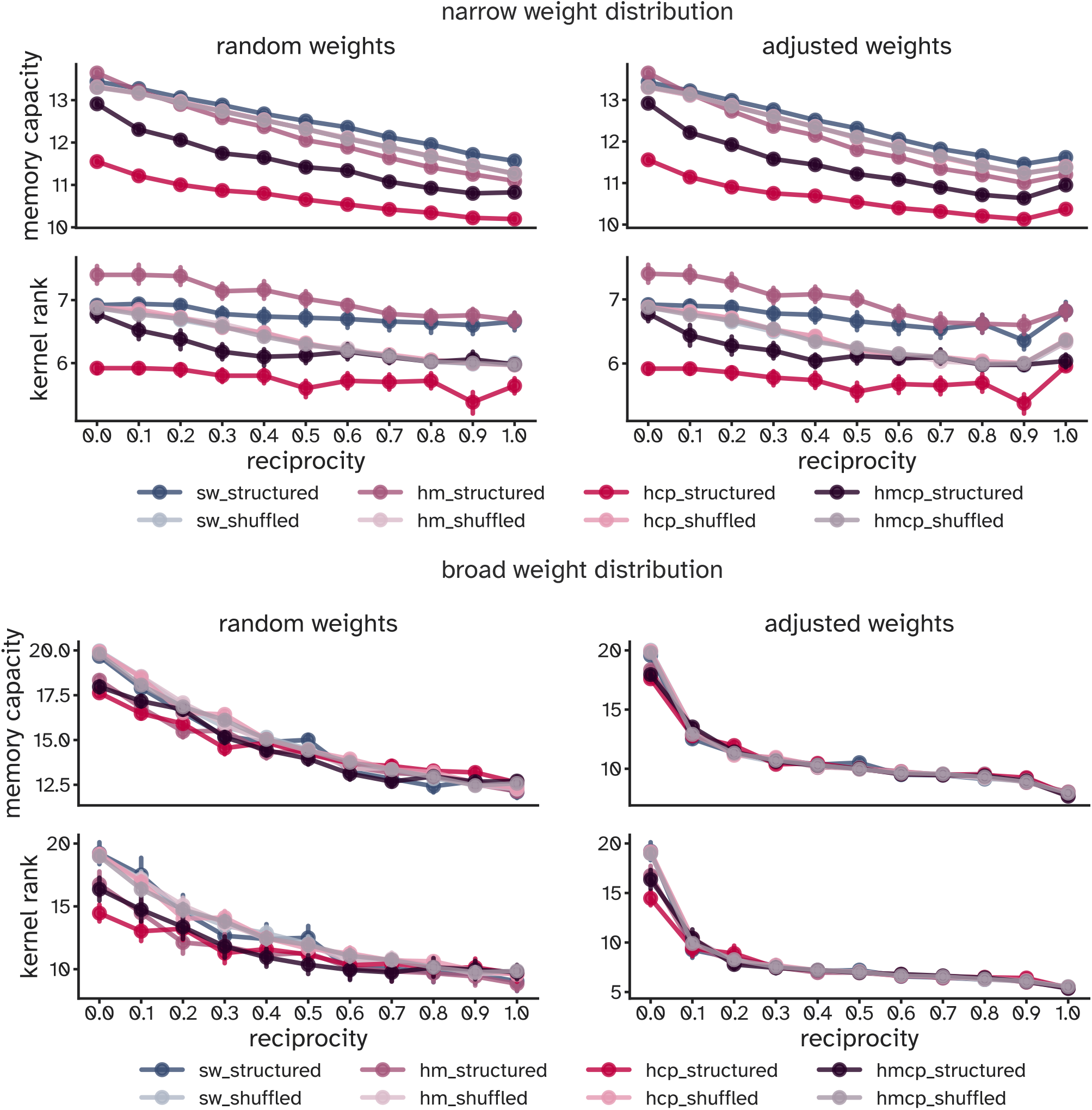
Effect of reciprocity on computation in large weighted networks (256 nodes). Memory capacity (MC) and kernel rank (KR) are shown for networks with log-normal weights (mean = 0.25) under random (left) and enforced (right) strength reciprocity. Results for low (σ = 0.1; top) and high (σ = 0.9; bottom) variability show that reciprocity impairs performance across all conditions, with more pronounced effects in longer-tailed weight distributions. Trends replicate findings in smaller networks. Error bars indicate standard deviation across 50 trials.

**Figure S7.**
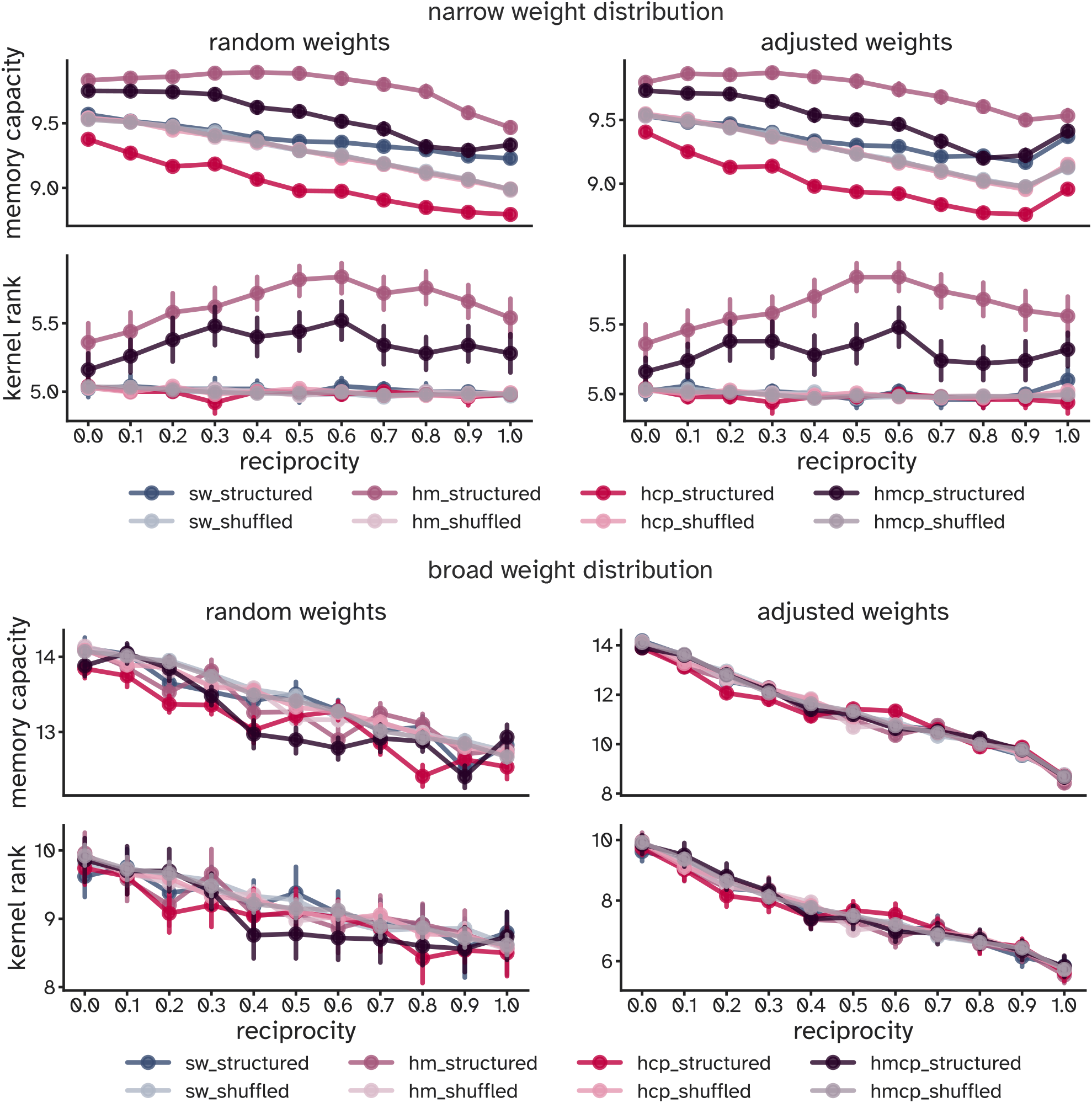
Impact of reciprocity on memory and kernel rank in weighted networks with random and adjusted strength reciprocity. Top and bottom panels show results for networks with weights sampled from log-normal distributions with standard deviations of 0.1 and 0.9, respectively (mean = 0.25 in both). Weights were assigned to binary network backbones generated under the higher connectivity regime, maintaining a mean out-degree of 10 while introducing log-normal heterogeneity. Across both weight regimes, increasing link reciprocity leads to systematic declines in MC and KR. Longer-tailed distributions (bottom panel) yield higher overall performance but exhibit sharper declines as reciprocity increases. Results shown for 64-node networks; error bars indicate standard deviation across 50 trials.

**Figure S8.**
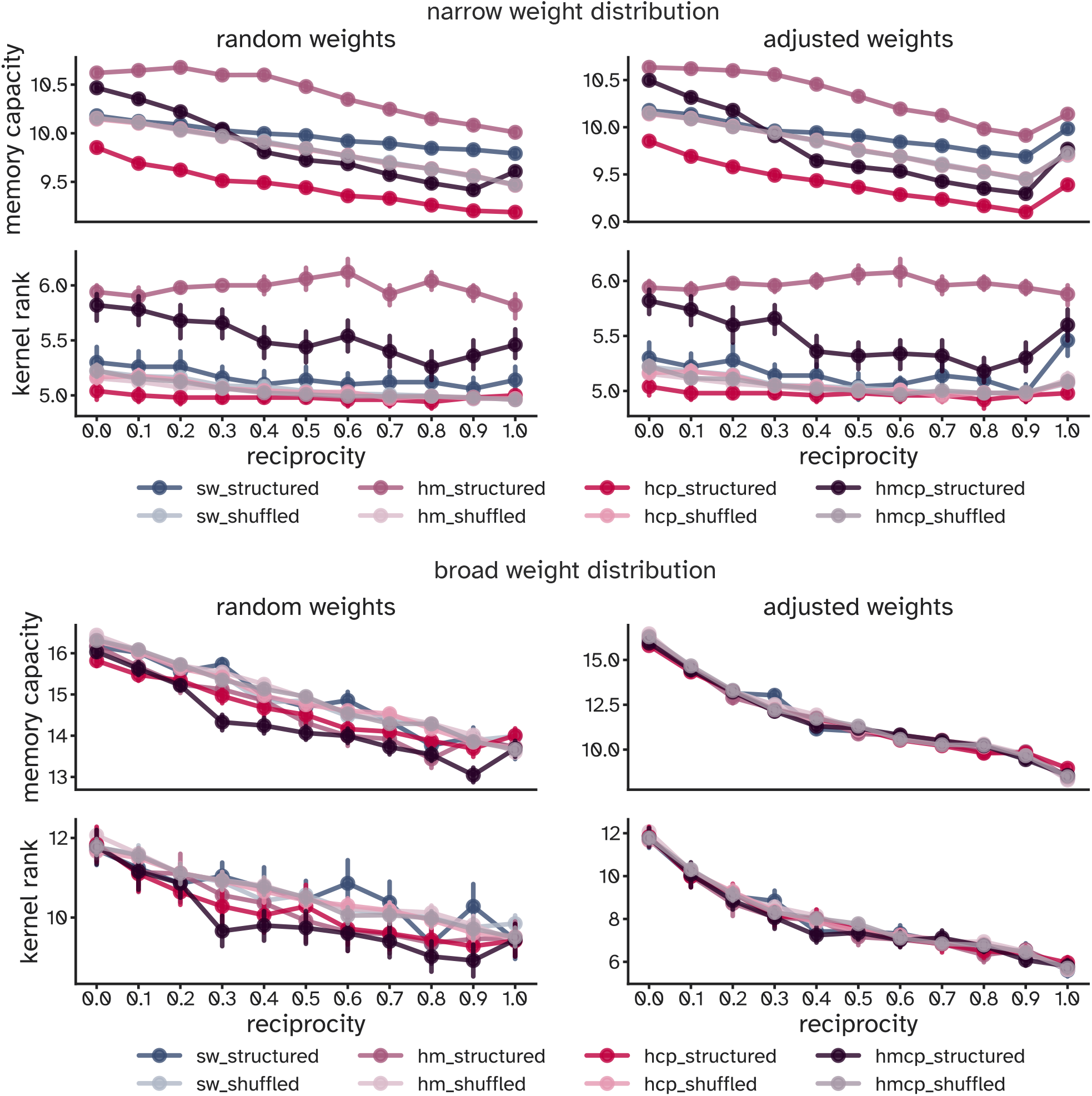
Impact of reciprocity on memory and kernel rank in weighted networks with random and adjusted strength reciprocity. Top and bottom panels show results for networks with weights sampled from log-normal distributions with standard deviations of 0.1 and 0.9, respectively (mean = 0.25 in both). Weights were assigned to binary network backbones generated under the higher connectivity regime, with mean out-degree fixed at 10. In all cases, increasing link reciprocity leads to consistent declines in MC and KR, particularly under longer-tailed weight distributions (bottom panel). Results shown for 128-node networks; error bars represent standard deviation across 50 network realizations.

**Figure S9.**
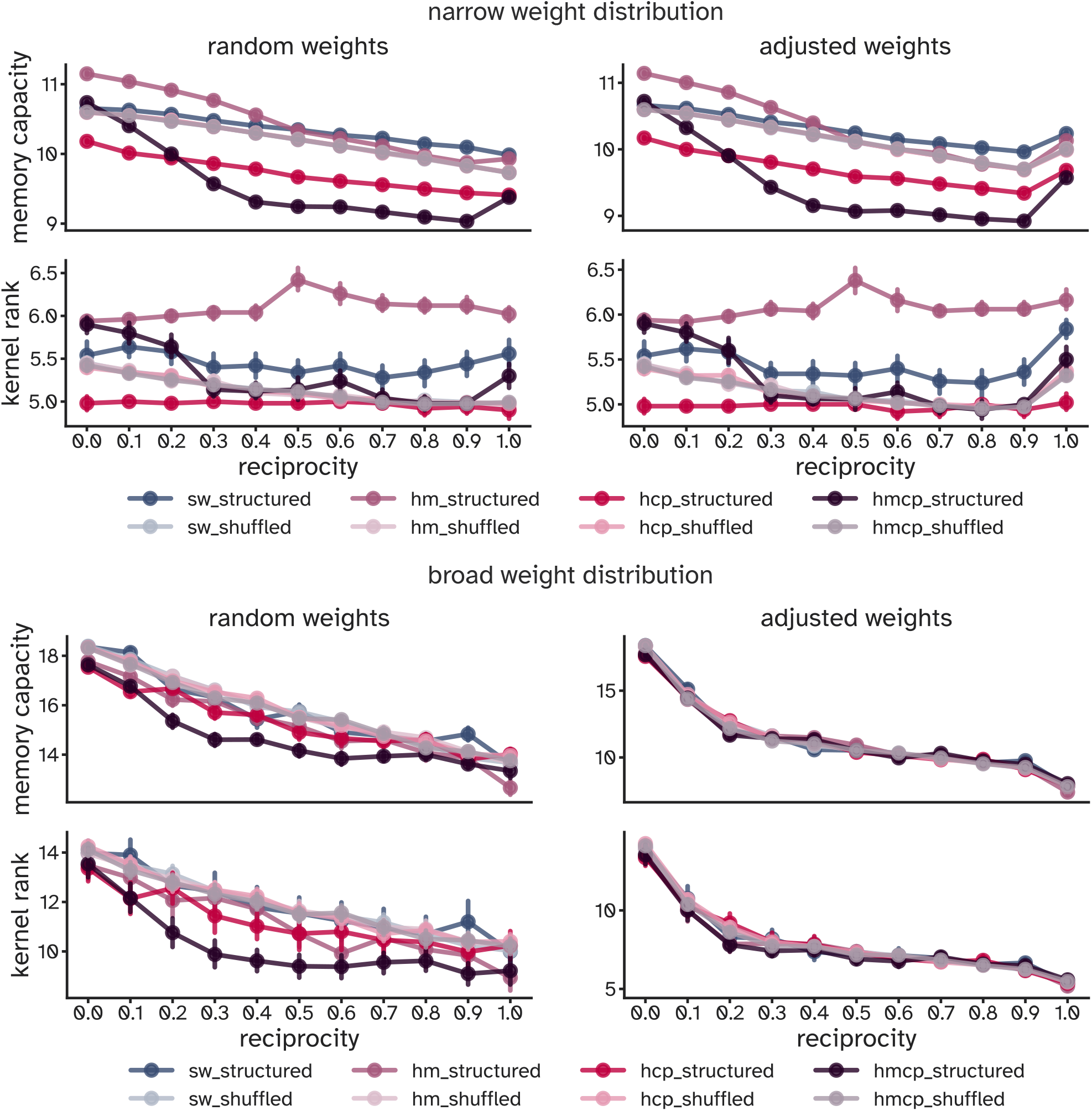
Impact of reciprocity on memory and kernel rank in weighted networks with random and adjusted strength reciprocity. Top and bottom panels show results for networks with weights drawn from log-normal distributions with standard deviations of 0.1 and 0.9, respectively (mean = 0.25 in both). All weights were applied to binary connectivity profiles generated in the higher connectivity regime, ensuring fixed out-degree (*k* = 10) and consistent density across architectures. Both metrics exhibit a steady decline with increasing reciprocity, and the effect is amplified for high-variance weights (bottom panel). Results are shown for 256-node networks; error bars indicate standard deviation across 50 repetitions.

**Figure S10.**
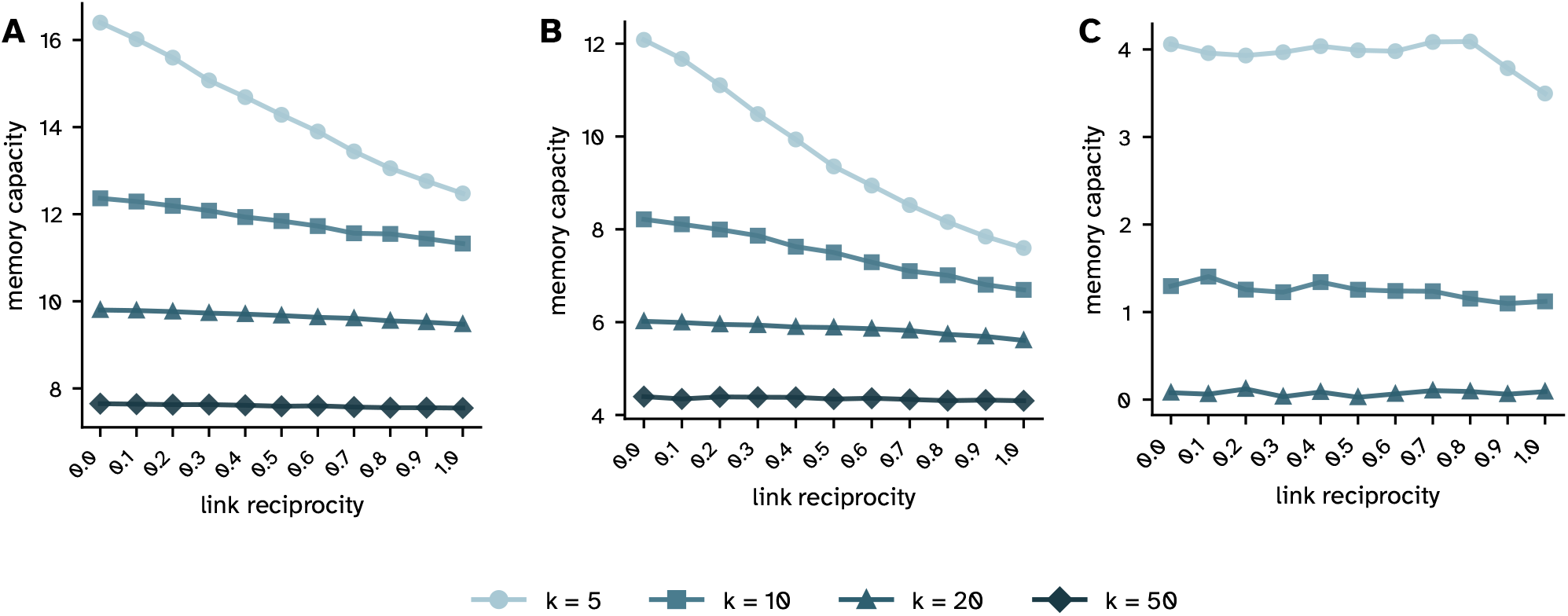
Memory capacity under varying connectivity, gain compensation, and without spectral normalization. All experiments used random binary networks with *N* = 400 nodes. Each data point represents the mean over 50 network realizations and 5 reservoir computing runs. Linear slopes reflect the change in MC across the full reciprocity range (0 → 1). **(A) Standard spectral normalization across connectivity levels**. MC declined linearly with reciprocity at all connectivity levels (all *R*^2^ ≥ 0.963, *P* < 0.001), with slopes steepening as density decreased: *k* = 5 (slope = −4.028), *k* = 10 (slope = −1.072), *k* = 20 (slope = −0.332), *k* = 50 (slope = −0.106). **(B) Input scaling by** *ρ* **to compensate for gain reduction**. When *W*_in_ was scaled by the raw spectral radius *ρ*, MC declines persisted across all connectivity levels (all *R*^2^ ≥ 0.708, *P* ≤0.001), with slopes of: *k* = 5 (slope = −4.681), *k* = 10 (slope = −1.602), *k* = 20 (slope = −0.379), *k* = 50 (slope = −0.082). At *k* = 5, 10, and 20, the *ρ*-scaled slopes were 1.16 ×, 1.49 ×, and 1.14 ×steeper than the normalized counterparts, confirming that gain compensation did not attenuate the reciprocity effect. At *k* = 50, the *ρ*-scaled slope was modestly shallower (0.78 ×; −0.082 vs −0.106), with a lower *R*^2^ (0.708 vs 0.963), suggesting that at high density the structural effect of reciprocity is weak and the linear fit less reliable in both conditions. **(C) Without spectral normalization**. Performance was strongly dependent on connectivity level. At *k* = 5, MC remained moderate across most reciprocity values (range: 3.50–4.09), with a notable decline at full reciprocity (4.09 ± 0.44 at *r* = 0.8 vs 3.50 ± 0.70 at *r* = 1.0), suggesting that the echo-state property is partially preserved at low density even without normalization. At *k* = 10, MC was substantially reduced (range: 1.10–1.41), indicating that the unnormalized spectral radius already impairs temporal memory. At *k* = 20, MC collapsed to near zero across all reciprocity levels (range: 0.03–0.12), reflecting a loss of the echo-state property. Networks with *k* = 50 produced MC = 0 at all reciprocity levels and are not shown.

**Figure S11.**
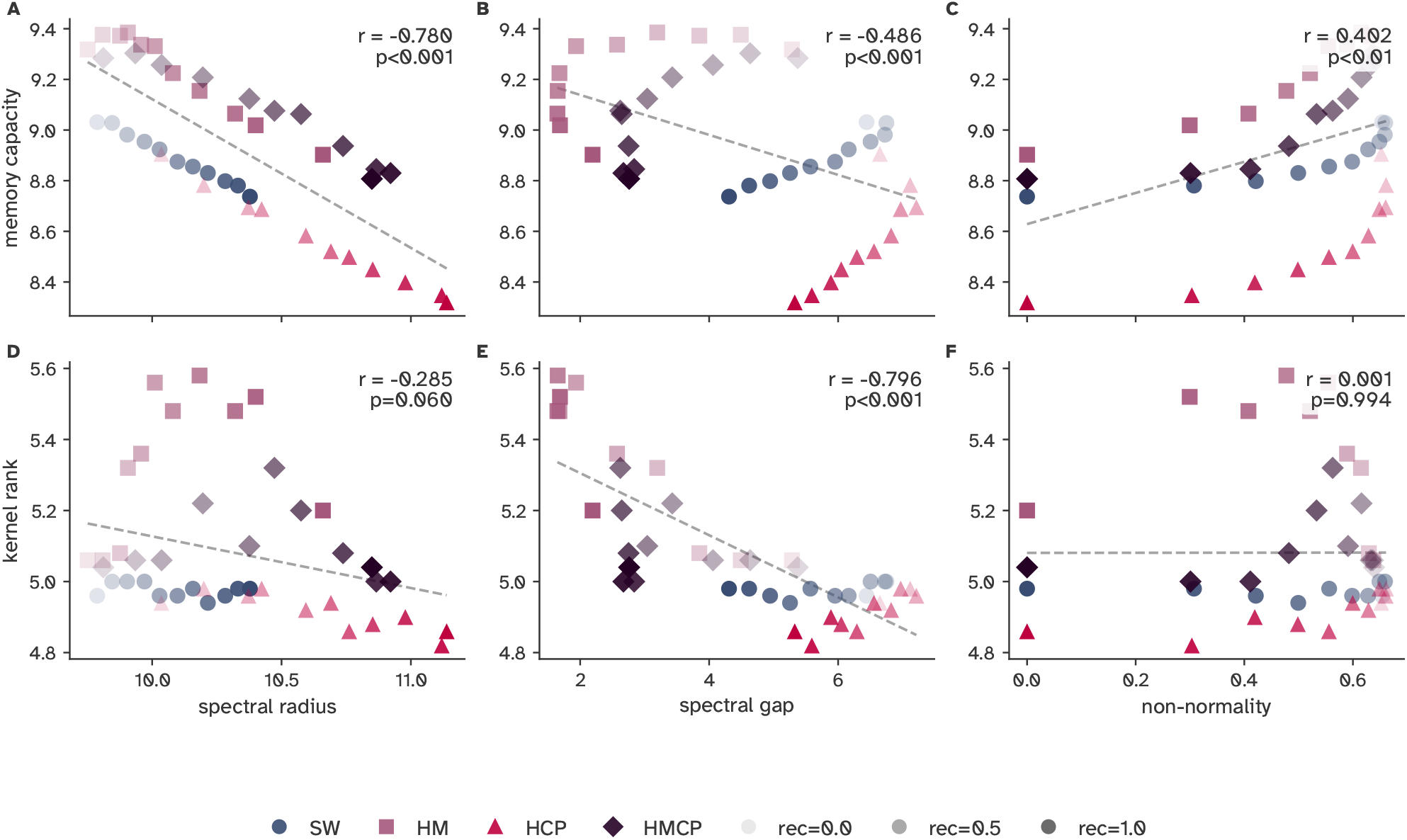
Spectral properties predict memory capacity and kernel rank across network architectures and reciprocity levels. Scatter plots show the relationship between three spectral properties (spectral radius, spectral gap, and non-normality) and two computational performance metrics (memory capacity, MC; kernel rank, KR) for binary networks with 64 nodes. Each point represents the mean over 50 network realisations for one (network type × reciprocity) combination. Network type is encoded by colour and marker shape (SW: circles; HM: squares; HCP: triangles; HMCP: diamonds); reciprocity is encoded by marker opacity (low reciprocity = faint, high reciprocity = opaque). Shuffled (random) counterparts are not shown, as their inclusion would obscure the topology-dependent patterns of interest. Dashed lines indicate linear regression fits across all structured networks combined. Pearson correlation coefficients (*r*) and *P*-values are reported for the pooled sample in each panel. For networks with higher connectivity (*k* = 10), spectral radius remained strongly negatively correlated with MC within each network type (SW *r* = −0.996, HM *r* = − 0.962, HCP *r* = −0.997, HMCP *r* = −0.976, all *P* < 0.001), though the pooled correlation was attenuated (*r* = −0.780, *P* < 0.001), indicating that denser connectivity narrows the range of variation across network types. Spectral gap showed divergent per-type correlations with MC (SW *r* = −0.962, HCP *r* = 0.839, HMCP *r* = 0.780, all *P* < 0.01; HM *r* = 0.594, *P* = 0.054) and a negative pooled correlation (*r* = − 0.486, *P* < 0.001), consistent with a Simpson’s paradox effect in which the pooling of heterogeneous network types reverses the apparent direction of the relationship. Non-normality was positively correlated with MC across all network types (SW *r* = 0.843, HM *r* = 0.942, HCP *r* = 0.781, HMCP *r* = 0.845, all *P* < 0.01; pooled: *r* = 0.402, *P* < 0.01), but showed no consistent relationship with KR at *k* = 10 (pooled: *r* = 0.001, *P* = 0.994). Spectral gap was strongly negatively correlated with KR at *k* = 10 (pooled: *r* = −0.796, *P* < 0.001), while spectral radius showed no significant relationship (pooled: *r* = −0.285, *P* = 0.060), suggesting that representational diversity is more sensitive to the gap structure of the spectrum than to its leading eigenvalue.

**Figure S12.**
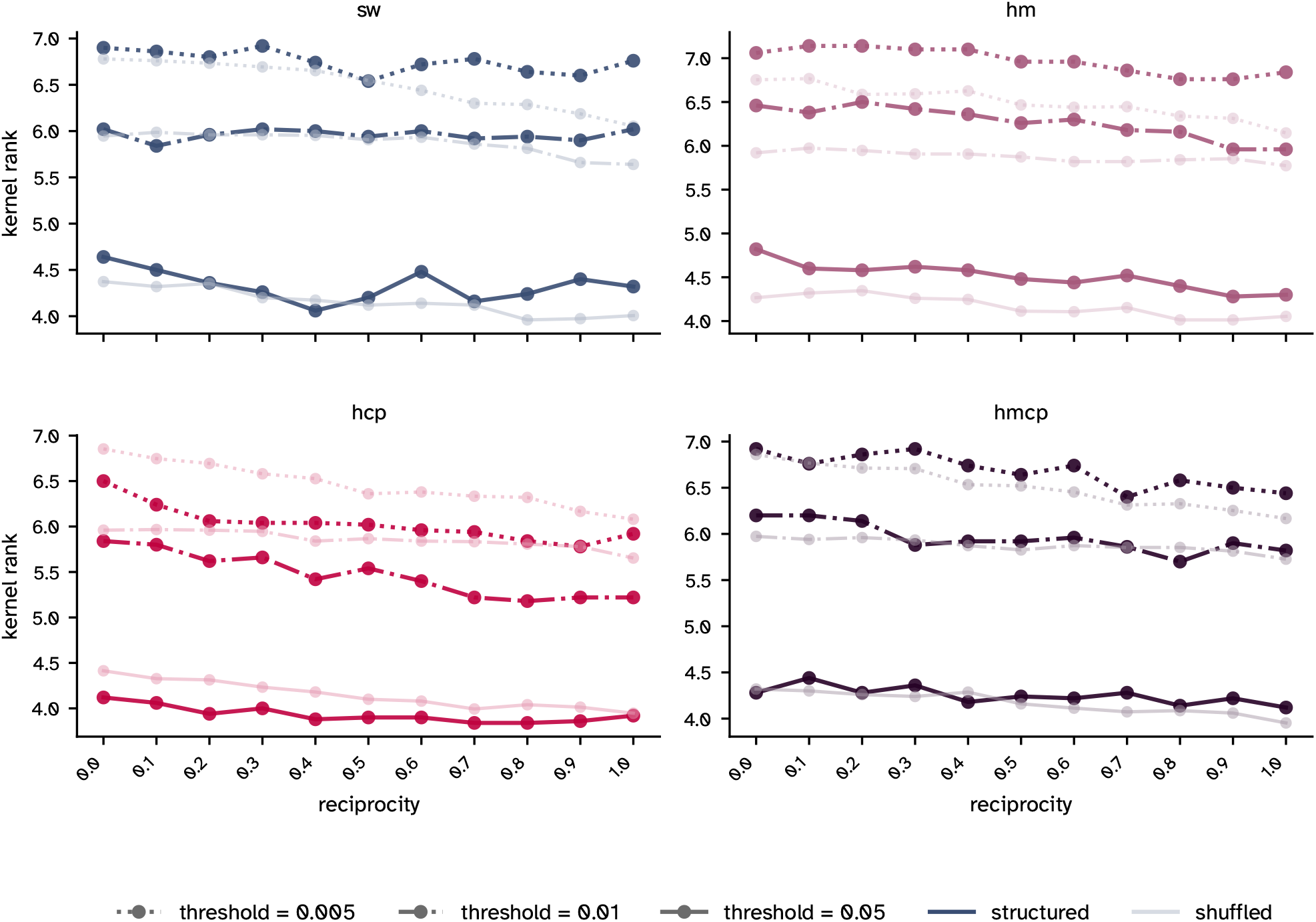
Kernel rank is robust to SVD threshold choice under binary connectivity. Kernel rank is shown as a function of link reciprocity for all four network architectures (SW, HM, HCP, HMCP) computed using three SVD thresholds (0.005, 0.01, 0.05), for both structured and random (null) networks. Results are shown for binary networks with *N* = 64 nodes and *k* = 6. Across all network types and thresholds, kernel rank declined consistently with increasing reciprocity. The absolute values of KR depend on the threshold, with smaller thresholds (e.g., 0.005_σ1_) retaining more singular values and yielding higher estimates, but the reciprocity-driven decline and the advantage of structured over shuffled networks are preserved across all threshold choices.

**Figure S13.**
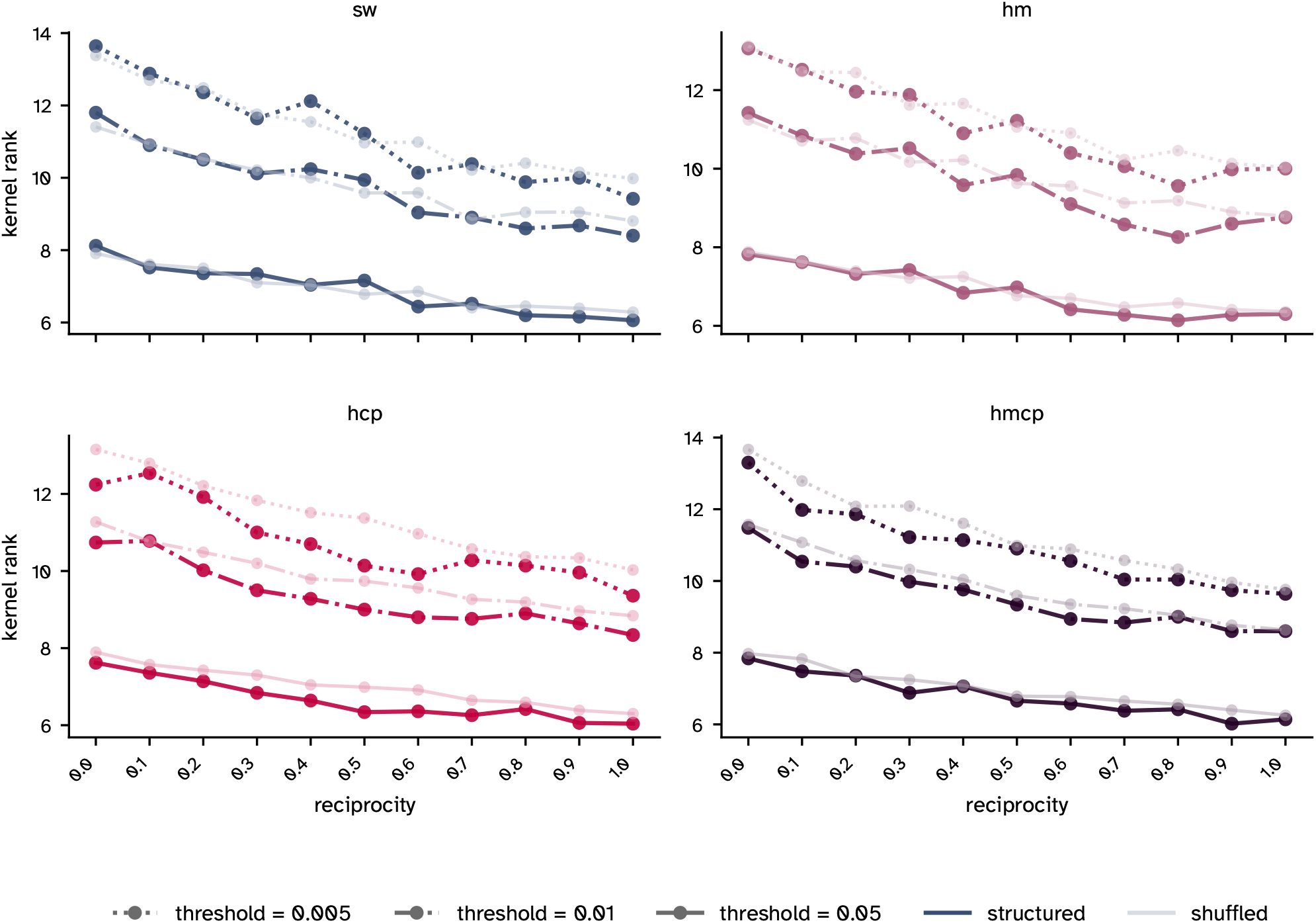
Kernel rank is robust to SVD threshold choice under random weight assignment. Same as the previous figure, but with log-normal weights (mean = 0.25, SD = 0.9) assigned randomly to binary network backbones (*N* = 64, *k* = 6). Across all network types and thresholds, kernel rank declined consistently with increasing reciprocity. The introduction of heterogeneous weights increased absolute KR values relative to the binary condition, but the reciprocity-driven decline remained consistent across all threshold choices.

**Figure S14.**
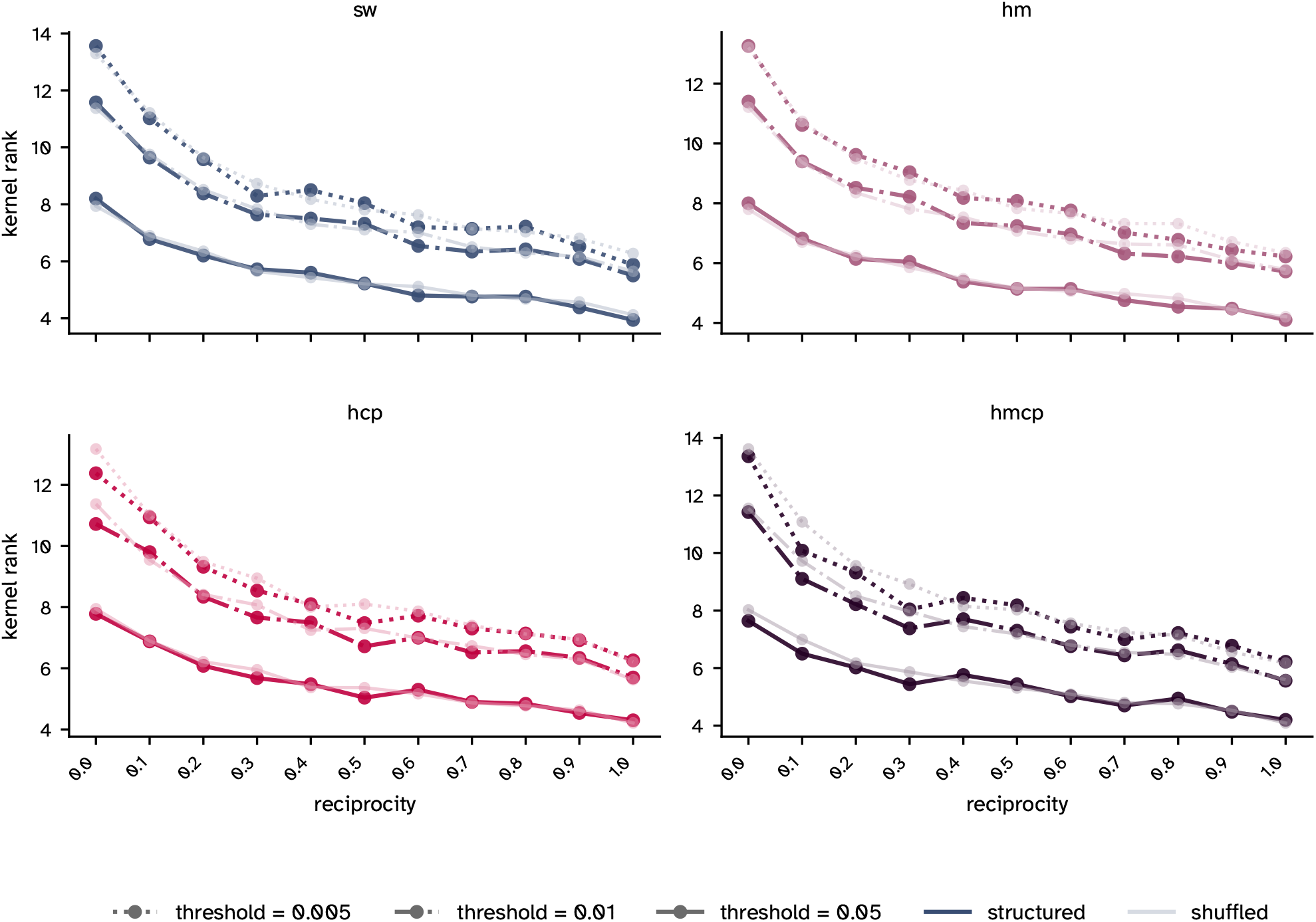
Kernel rank is robust to SVD threshold choice under enforced strength reciprocity. Same experimental setup as the previous figure, but with strength reciprocity explicitly aligned with link reciprocity using the NRC algorithm. Enforcing strength reciprocity amplified the decline in kernel rank across all network types and thresholds, consistent with the main text results. The qualitative pattern was consistent across all threshold choices, confirming that the KR results reported in the main text are not an artifact of the chosen threshold of 0.01_σ1_.

**Figure S15.**
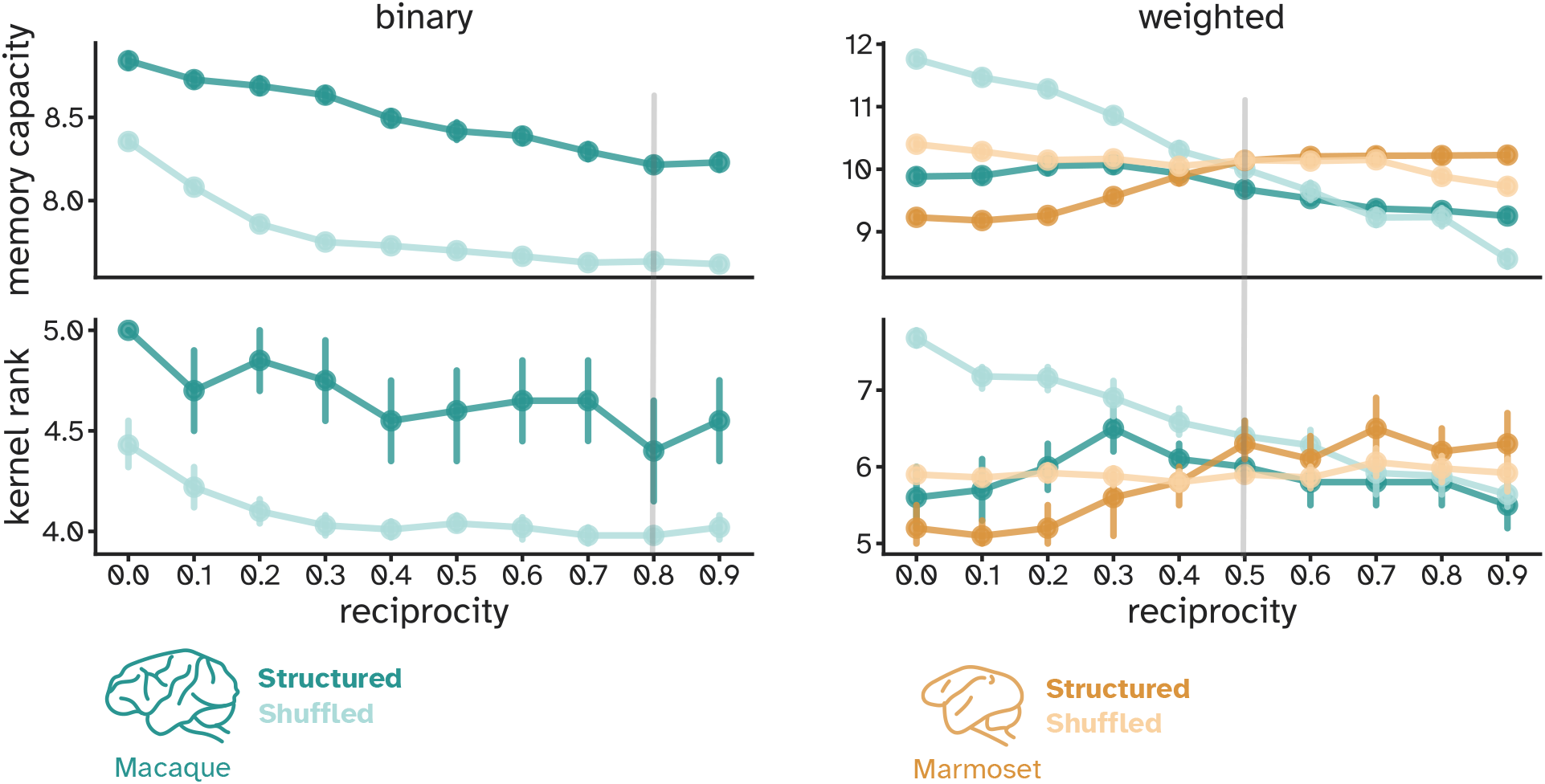
Increasing reciprocity alters memory capacity and kernel rank in binary and weighted empirical connectomes. Left panels show memory capacity and kernel rank as functions of link reciprocity for the binary macaque long-distance pathways connectome with 71 nodes and its degree-preserving rewired null models. At the original reciprocity value (0.8), the connectome-based reservoir model achieved a memory capacity of 8.23 ±0.23, whereas the rewired model, characterized by a lower link reciprocity value (∼0.25), reached a capacity of only 7.77 ± 0.19. This difference was highly significant (*P* < 0.001, Cohen’s *d* = 2.34). Both metrics decline with increasing reciprocity, with the empirical connectome outperforming null models across all reciprocity values. Right panels show memory capacity and kernel rank as functions of strength reciprocity for the weighted macaque visual cortex and marmoset cortical connectomes and their degree-preserving rewired null models. Reciprocity was adjusted in 20 trials for each network, with the spectral radius optimized to maximize memory capacity. Without normalization, increases in strength reciprocity lead to weight redistribution, which can artificially boost memory capacity and kernel rank in both empirical and null models.

**Figure S16.**
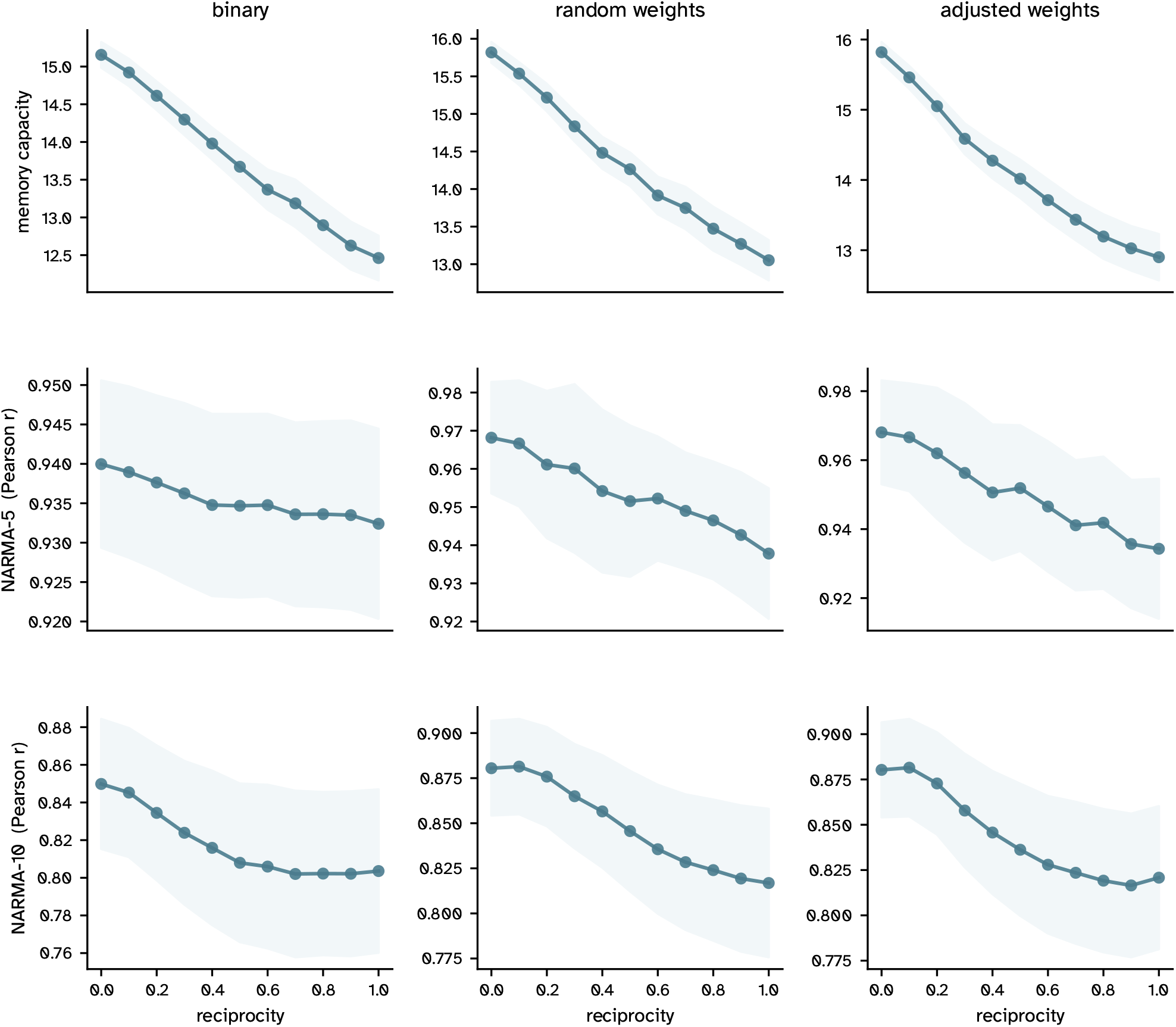
Reciprocity-driven decline generalizes to nonlinear tasks under narrow log-normal weights (SD = 0.1, mean = 0.25). Results are shown for random networks with *N* = 400 nodes and *k* = 6. Rows show memory capacity (MC), NARMA-5 Pearson correlation, and MA-10 Pearson correlation. Columns correspond to binary, random weight, and adjusted (enforced strength reciprocity) weight conditions. Shaded regions indicate mean ± SD across 50 network realizations. Under binary connectivity, MC declined linearly (slope = < 2.78, *R*^2^ = 0.996, *P* < 0.001; 15.16 ± 0.19 to 12.46 ± 0.29), while NARMA-5 showed a shallower but significant decline (slope = −0.007, *R*^2^ = 0.907, *P* < 0.001; 0.940 ± 0.011 to 0.932 ± 0.012). NARMA-10 exhibited a nonlinear profile: steeper at low reciprocity and flattening beyond *r* ≈0.5. This trend is reflected in a lower linear fit quality (slope = − 0.050, *R*^2^ = 0.853, *P* < 0.001; 0.850 ± 0.036 to 0.804 ± 0.046). Under random weight assignment (SD = 0.1, mean = 0.25), declines were more pronounced across all metrics (MC: slope = −2.81, *R*^2^ = 0.991, *P* < 0.001, 15.82 ± 0.12 to 13.05 ± 0.25; NARMA-5: slope = − 0.029, *R*^2^ = 0.976, *P* < 0.001, 0.968 ± 0.013 to 0.938 ±0.017; NARMA-10: slope = −0.074, *R*^2^ = 0.974, *P* < 0.001, 0.881 ± 0.026 to 0.817 ± 0.041), with narrow variance bands reflecting stable performance across network instances. Under enforced strength reciprocity (adjusted weights), declines were comparable in magnitude to the random condition for this weight distribution, with linear and exponential fits yielding similar goodness of fit (MC: decay rate = −0.21, *R*^2^ = 0.986, 15.82 ± 0.14 to 12.90 ± 0.33; NARMA-5: decay rate = −0.037, *R*^2^ = 0.978, 0.968 ± 0.013 to 0.934 ± 0.021; NARMA-10: = −0.087, *R*^2^ = 0.925, 0.880 ± 0.027 to 0.821 ± 0.039).

**Figure S17.**
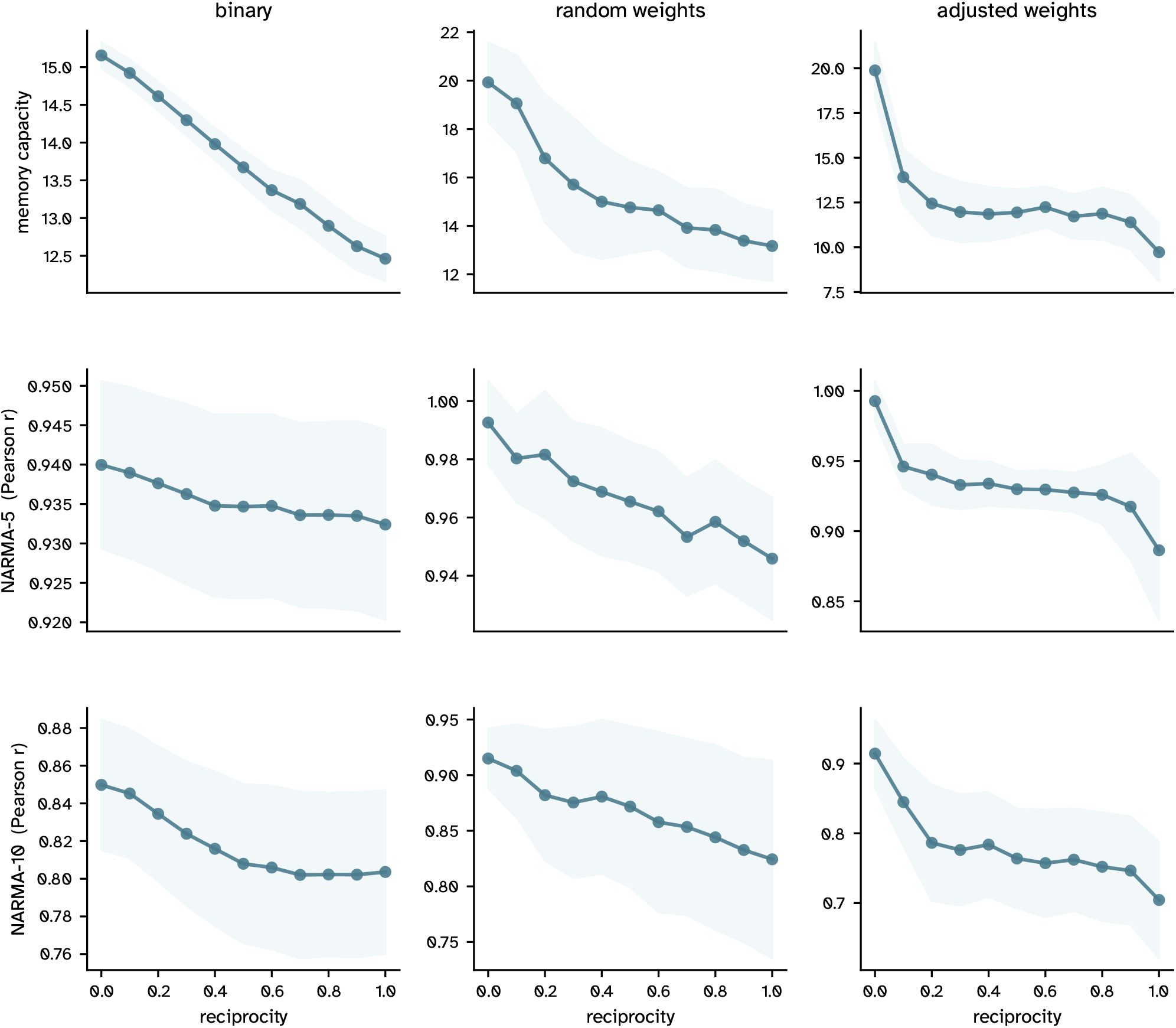
Reciprocity-driven decline generalizes to nonlinear tasks under broad log-normal weights (SD = 0.9, mean = 0.25). Same experimental setup as Supplementary Figure S16, with weights drawn from a broader log-normal distribution. The binary column is identical to Supplementary Figure S16. Shaded regions indicate mean ± SD across 50 network realizations. Under random weight assignment (SD = 0.9), broader weights yielded substantially higher baseline performance but steeper reciprocity-driven declines, with wider variance bands reflecting greater across-instance variability (MC: slope = −6.30, *R*^2^ = 0.861, *P* < 0.001, 19.93 ± 1.08 to 13.17 ±1.11; NARMA-5: slope = −0.042, *R*^2^ = 0.950, *P* < 0.001, 0.993 ± 0.004 to 0.946 ± 0.019; NARMA-10: slope = − 0.084, *R*^2^ = 0.964, *P* < 0.001, 0.915 ± 0.022 to 0.824 ± 0.084). Under enforced strength reciprocity (adjusted weights), performance declined most steeply, with an exponential decay profile most evident in MC and NARMA-10. The lower goodness-of-fit values (*R*^2^ = 0.569–0.764) reflect high across-trial variability rather than absence of a trend, which is visually clear in all panels. Endpoint values confirm the severity of the decline: MC dropped from 19.88 ±1.14 to 9.72 ±1.43 (decay rate = −0.485); NARMA-5 from 0.993 ±0.004 to 0.886 ±0.065 (decay rate = −0.069); and NARMA-10 from 0.914 ± 0.022 to 0.704 ± 0.080 (decay rate = −0.190). Together with Supplementary Figure S16, these results confirm that enforced strength reciprocity imposes a consistent computational cost across tasks, and that broader weight distributions amplify rather than buffer this effect.

**Figure S18.**
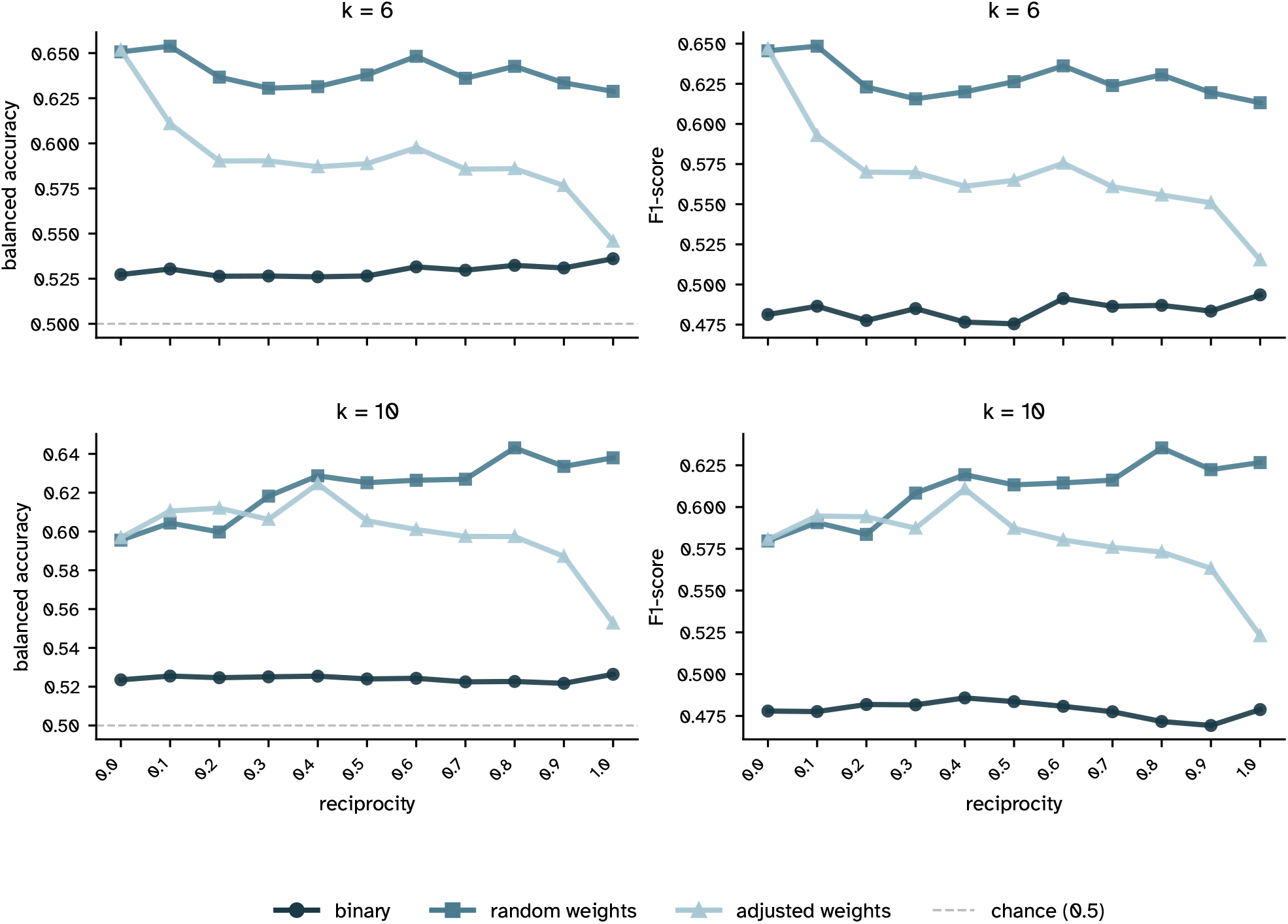
Reciprocity impairs context-dependent classification performance in weighted networks. Balanced accuracy (left) and macro F1 score (right) on the Mante context-dependent decision-making task (Mante et al., 2013) are shown as functions of reciprocity for binary, random weight, and adjusted (enforced strength reciprocity) conditions, at *k* = 6 (top) and *k* = 10 (bottom). Networks had *N* = 400 nodes. The dashed line indicates chance-level performance (balanced accuracy = 0.5). Under binary connectivity, performance remained near chance across all reciprocity levels at both connectivity regimes. Under random weight assignment at *k* = 6, performance declined modestly with increasing reciprocity (balanced accuracy: 0.651 ±0.018 at reciprocity = 0 to 0.629 ± 0.058 at reciprocity = 1.0; macro F1: 0.646 ± 0.021 to 0.613 ± 0.072). Under enforced strength reciprocity at *k* = 6, the decline was more pronounced (balanced accuracy: 0.652 ±0.018 to 0.546 ± 0.030; macro F1: 0.647 ± 0.021 to 0.515 ±0.043), consistent with the amplifying effect of strength reciprocity observed across all other tasks. At *k* = 10, random weight assignment showed no consistent decline, while enforced strength reciprocity produced a moderate reduction in both metrics (balanced accuracy: 0.597 ±0.022 to 0.553 ±0.034; macro F1: 0.581 ± 0.030 to 0.523 ± 0.049), suggesting that the effect of reciprocity on classification is attenuated at higher connectivity levels. Complete mean and standard deviation values across all reciprocity levels and conditions are provided in Supplementary Table S3.

**Figure S19.**
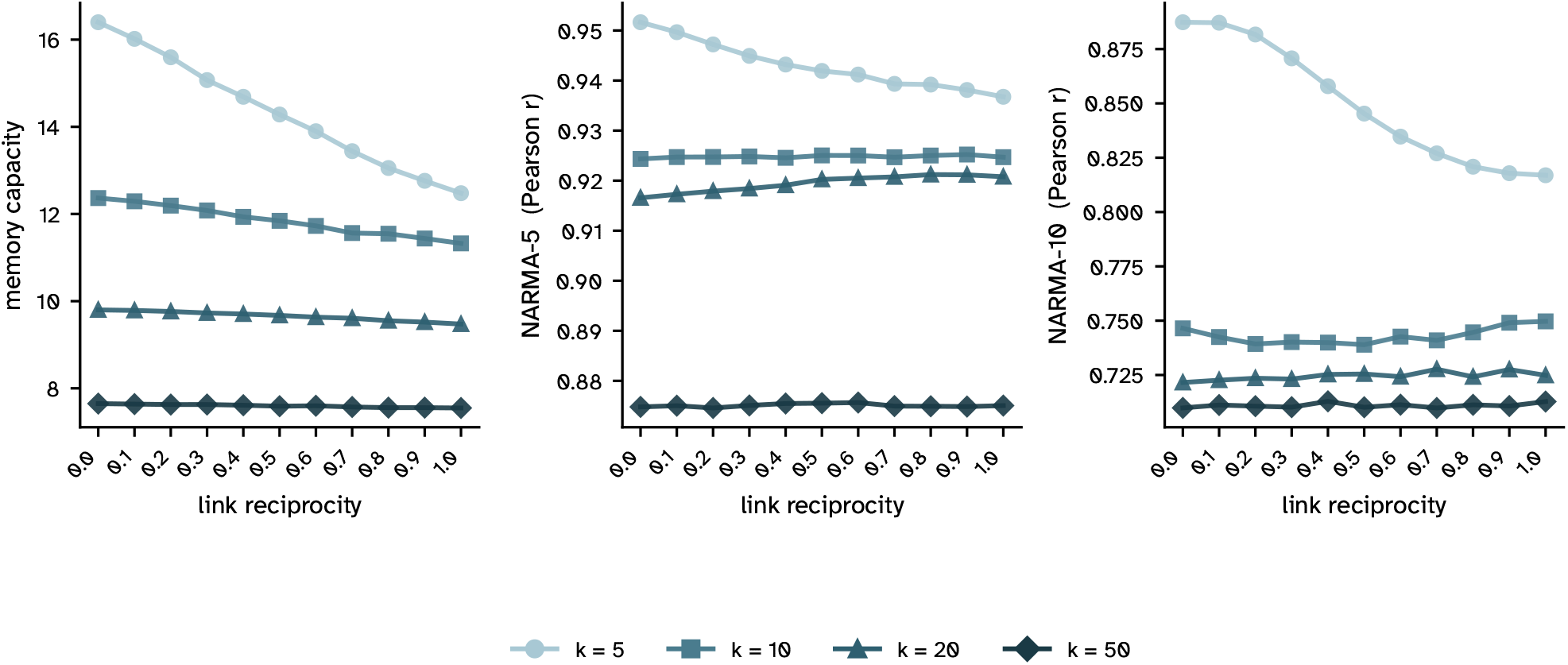
Effect of out-degree on the relationship between link reciprocity and computational performance. Shown are memory capacity (MC; left), NARMA-5 Pearson correlation (centre), and NARMA-10 Pearson correlation (right) as functions of link reciprocity for binary random networks with *N* = 400 nodes and four mean out-degrees: *k* = 5, 10, 20, and 50. Each data point represents the mean across 50 independently generated network instances; error bars are omitted for clarity. Higher out-degree was associated with lower performance across all three tasks and all reciprocity levels. For MC, increasing link reciprocity produced a consistent and near-perfectly linear decline at all out-degrees (linear regression, all *P* < 0.001; *k* = 5: slope = − 4.03, *R*^2^ = 0.997; *k* = 10: slope = − 1.07, *R*^2^ = 0.994; *k* = 20: slope = −0.33, *R*^2^ = 0.985; *k* = 50: slope = −0.11, *R*^2^ = 0.963). The absolute magnitude of the decline diminished with increasing density, ranging from 16.40 ±0.20 to 12.48 ± 0.36 at *k* = 5, and from 7.65 ±0.033 to 7.55 ± 0.046 at *k* = 50 (mean SD across 50 trials, at reciprocity = 0 and 1, respectively). For NARMA-5 and NARMA-10, the reciprocity effect was out-degree-dependent. Only the sparsest condition (*k* = 5) exhibited a significant negative trend (NARMA-5: slope = −0.014, *R*^2^ = 0.961, *P* < 0.001; NARMA-10: slope = −0.084, *R*^2^ = 0.963, *P* < 0.001), with Pearson *r* declining from 0.952 ±0.010 to 0.937 ± 0.012 and from 0.887 ± 0.025 to 0.817 ± 0.041, respectively. At *k* = 10 and *k* = 50, trends were negligible and non-significant in both NARMA tasks (all *P* > 0.06), with endpoint values differing by less than 0.001 in both cases. At *k* = 20, a small positive trend was observed (NARMA-5: slope = +0.005, *R*^2^ = 0.903, *P* < 0.001; NARMA-10: slope = +0.004, *R*^2^ = 0.536, *P* = 0.010), indicating a marginal improvement in nonlinear task performance with increasing reciprocity in moderately dense networks. Complete mean and standard deviation values across all out-degrees and reciprocity levels are provided in Supplementary Table S4.

**Supplementary Table S1 | Memory capacity (MC) and kernel rank (KR) of structured and null/random networks across all experimental conditions**. Each row corresponds to one combination of network type (SW, HM, HCP, HMCP), network size (*N*), out-degree (*k*), weight regime (binary; random narrow/broad, σ = 0.1 /0.9; adjusted narrow/broad, σ = 0.1 /0.9), and reciprocity level (0–1 in steps of 0.1). For each condition, mean and standard deviation of MC and KR are reported separately for structured and null/random networks. Null/random networks are weight-shuffled surrogates that preserve mean out-degree and weight distribution but destroy structured connectivity. *P*-values reflect pairwise comparisons between structured and null/random networks at each reciprocity level (two-sided *t*-test; Welch’s *t*-test for *N* = 1024). Cohen’s *d* quantifies the effect size of the structured vs. null/random difference. For *N* = 1024, experiments were conducted with 30 trials and 2 null/random networks (vs. 50 trials and 5 null/random networks for *N* ≤512). NaN entries in KR statistics indicate conditions where both structured and null/random networks produced zero-variance KR values, rendering statistical comparison undefined. *P* < 0.0001 is reported as such rather than 0.

**Supplementary Table S2 | Trend analysis of memory capacity (MC) and kernel rank (KR) across reciprocity levels for all experimental conditions**. Each row corresponds to one combination of network type (SW, HM, HCP, HMCP), network size (*N*), and out-degree (*k*). For binary, random, and adjusted-narrow weight regimes, a linear model (*y* = *a* · *x* + *b*) was fitted to the mean MC or KR of structured networks as a function of reciprocity (0 →1); the slope reflects the change in performance per unit reciprocity. For the adjusted-broad weight regime, an exponential model (*y* = *a* · *e*^*b*·*x*^ +*c*) was fitted; the decay rate *b* is reported in the dedicated decay rate column and is otherwise NaN. All slopes and decay rates are expressed per unit reciprocity (scale 0 →1). The *P*-value of the slope (or decay rate) is derived from linear regression for linear models, and approximated from the standard error o*b* via a *t*-test for exponential models. *R*^2^ quantifies goodness of fit. For *N* = 1024, experiments were conducted with 30 trials and 2 null/random networks (vs. 50 trials and 5 null/random networks for *N* ≤512); Welch’s *t*-test was used for KR comparisons. NaN entries in KR slope and *R*^2^ indicate conditions where KR values showed no variation across trials or reciprocity levels, rendering trend fitting undefined.

**Supplementary Table S3 | Context-dependent classification performance across reciprocity levels, connectivity regimes, and weight conditions**. Mean and standard deviation of balanced accuracy and macro F1 score on the Mante context-dependent decision-making task (Mante et al., 2013) for networks with *N* = 400 nodes, at out-degree *k* = 6 and *k* = 10. Results are shown for three weight conditions (binary, random weight, and enforced strength reciprocity) across link reciprocity values from 0 to 1.0.

**Supplementary Table S4 | Effect of out-degree on the relationship between link reciprocity and computational performance**. Mean and standard deviation of memory capacity (MC), NARMA-5 Pearson correlation, and NARMA-10 Pearson correlation for binary random networks with *N* = 400 nodes across four mean out-degrees (*k* = 5, 10, 20, 50) and link reciprocity levels from 0 to 1.0 in steps of 0.1. Each value is averaged across 50 independently generated network instances.

